# Differential proteomics of bacteria grown *in vitro* and *in planta* reveals functions used during growth on maize roots

**DOI:** 10.1101/2025.06.02.657423

**Authors:** Anna-Katharina Garrell, John Cheadle, Nathan Crook, Gaurav Pal, Alecia N. Septer, Maggie R. Wagner, Ashley E. Beck, Manuel Kleiner

## Abstract

Microbes are ubiquitous in the rhizosphere and play crucial roles in plant health, yet the metabolisms and physiologies of individual species *in planta* remain poorly understood. In this study, we examined microbial gene expression in response to the maize root environment for seven bacterial species originally isolated from maize roots. We grew each species individually, both *in vitro* in a minimal medium and *in planta*, and used differential proteomics to identify functions upregulated specifically when bacteria are grown on maize roots. We identified between 1,500 and 2,100 proteins from each species, with 20-60% of these proteins being differentially abundant between the two conditions. While we found that transporter proteins were upregulated in all species *in planta*, all other differentially abundant functions varied greatly between species, suggesting niche specialization in root-associated microbes. Indeed, *in vitro* assays confirmed that *Curtobacterium pusillum* likely degrades plant hemicellulose, *Enterobacter ludwigii* may benefit the plant by phosphate solubilization, and *Herbaspirillum robiniae* colonizes maize roots more effectively when both of its Type VI Secretion Systems are functional. Together, our findings highlight both conserved and species-specific bacterial strategies for growth in the root environment and lay a foundation for future work investigating the mechanisms underlying plant-microbiota interactions.

**Importance:** Bacteria that live on and around plant roots are important for plant growth and health, yet we still know relatively little about how individual bacterial species behave in this environment. In this study, we looked at seven bacterial species originally isolated from maize roots to understand how they change their metabolism and physiology when grown on the plant versus when grown under laboratory conditions. By doing this, we identified key strategies bacteria use to survive and thrive in the root environment, including changes in nutrient uptake, metabolism, and secretion systems. We also substantiated some of these behaviors using lab experiments and bacterial mutants. Understanding these species-specific functions helps us learn how bacteria establish themselves on roots and interact with the plant. This knowledge is critical for future efforts to design effective microbial communities that improve crop performance and resilience.

## Introduction

Plant-associated bacteria represent an important component of agricultural ecosystems, as they are ubiquitous and have been shown to support plant health and growth (1–4). As agricultural production faces increasing challenges from climate change, there is a vested interest in harnessing beneficial bacteria to improve crop resilience and productivity. While the application of microbial inoculants has emerged as a promising strategy, realizing their full potential requires a deeper understanding of microbial colonization, function, and adaptation in plant environments. As such, there is an increasing push to not only identify which microbes are present and their functional potential, but to also understand which microbial functions are actively expressed in their respective environments. This will ultimately lead to a better understanding of the mechanisms underlying community assembly, microbial colonization and persistence on plant roots, and beneficial plant-microbe interactions.

In an effort to identify bacterial functions that affect plant health, numerous studies have used transposon mutagenesis, transcriptomics, and other ‘omics approaches to probe plant-microbe interactions (5–14). These studies have revealed that phosphate uptake, nitrogen metabolism, and amino acid transport and metabolism, among many others, may contribute to microbial colonization and survival in the root environment. While these studies provided insights into functions of relevance for plant-associated microorganisms, our understanding of which functions are important for plant-associated as compared to free-living growth is limited. Additionally, while hundreds of microbial species are known to colonize plants and the rhizosphere (15–18), gene expression *in planta* has been measured for few; thus, our understanding of which functions are of general relevance for plant-associated growth, versus which functions are specific to particular microbial taxa, is limited. To address these knowledge gaps, we studied gene expression of protein-coding genes in seven microbial species grown individually *in vitro* and *in planta* using proteomics.

Maize is the second most widely grown crop in the world serving as a main source for human food, livestock feed, and biofuel (19). Maize is also a key model system for plant genetics (20, 21), and more recently, for plant-microbe interactions (22–24). In 2017, Niu *et al.* developed a simplified and representative synthetic community (SynCom) for maize containing seven bacterial species isolated from maize roots: *Brucella pituitosa*, *Chryseobacterium indologenes*, *Curtobacterium pusillum*, *Enterobacter ludwigii*, *Herbaspirillum robiniae*, *Pseudomonas putida*, and *Stenotrophomonas maltophilia* (24). These isolates were found to consistently colonize maize roots for up to 15 days and have since been used to investigate microbe-dependent heterosis (23) and metabolic microbe-microbe interactions (25). Additional tools and methods have also been developed for this community (26–28), making it an excellent model for studying plant-microbe interactions.

Proteomics allows for the identification and quantification of thousands of proteins in host-associated microbes, providing insight into the functions driving microbial phenotypes and interactions (29–31). With the development and advancements made in the field of proteomics and metaproteomics over the last two decades, our ability to identify the proteins responsible for phenotypes in complex systems has improved dramatically and deepened our understanding of microbe-microbe and microbe-plant interactions (12, 32–34). Differential proteomics, in particular, allows us to probe how changes in the biotic and abiotic environment affect gene expression of both microbes and hosts.

Dissecting the functions underlying interactions between plant roots and individual bacterial species is essential to understanding the specific traits that enable root colonization and persistence. Therefore, we focused on single-species interactions with maize roots by growing seven maize-root bacterial isolates individually *in vitro* in a defined, minimal medium and *in planta* on sterile maize roots (24). We used proteomics to identify bacterial proteins that were differentially abundant between the two conditions, allowing us to determine metabolic pathways and physiological functions that are upregulated during growth *in planta* and thus potentially important for growth in or on roots. We then used *in vitro* assays to further investigate hemicellulose degradation in *C. pusillum* and *H. robiniae* and phosphorus solubilization in all seven species, and generated functional knockouts of the Type VI Secretion System (T6SS) in *H. robiniae* to investigate its role in plant-microbe interactions.

## Results and Discussion

### Hundreds to over a thousand proteins were differentially abundant in response to growth on maize roots

To determine bacterial genes relevant for growth on maize roots, we grew seven bacterial species individually on sterile maize roots (*in planta*) and in a defined, minimal medium (*in vitro*). We used LC-MS/MS-based proteomics to quantify relative protein abundances in each condition. This differential proteomics approach allowed us to identify bacterial genes that are upregulated in the maize root environment. Using this method, we identified between 1,507 and 2,159 bacterial proteins for each bacterial species (Table 1). Approximately 20-60% of these proteins were differentially abundant (Student’s T-Test, Benjamini-Hochberg FDR corrected p-value (*q*) < 0.05) between conditions, indicating environment-specific metabolic and physiological responses in each of the species.

**Table 1.**
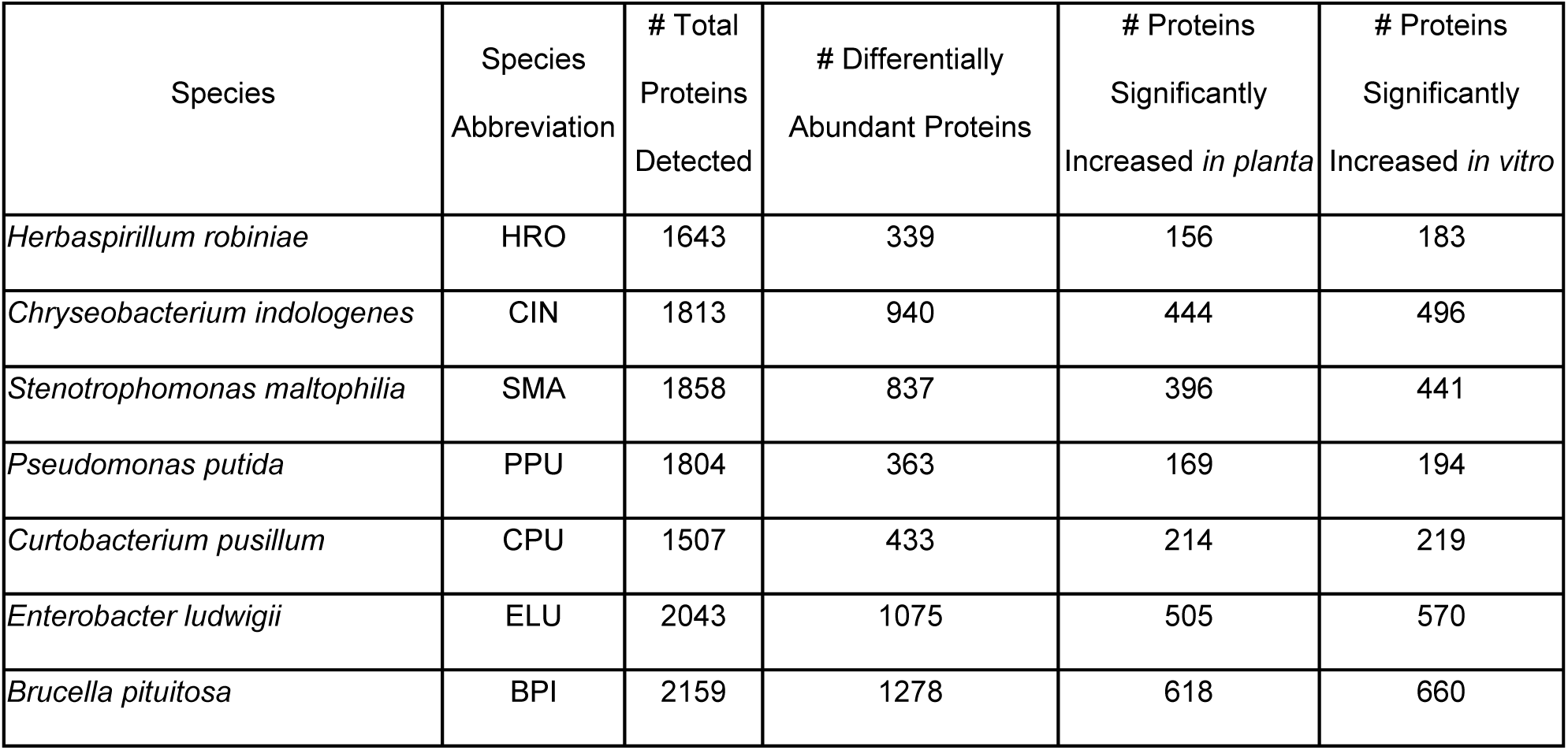
Overview of total detected and differentially abundant proteins (Student’s T-Test, Benjamini-Hochberg corrected p-value (*q*) < 0.05) in seven bacterial species grown *in vitro* and *in planta*.

To identify the functional roles of the differentially abundant proteins, we annotated these proteins, assigning a total of 28 broad categories and 133 sub-categories (Supplementary Data 1). It is important to note that included in the broad categories is a category called “Unknown Function”. This category contained proteins for which we attempted to assign a function but were unable to find a confident match.

Of the ∼3,000 differentially abundant proteins we were able to annotate, most belonged to three primary functional categories: transport, gene expression, and carbon metabolism (Fig. 1A). The extent and condition in which these and other metabolic and physiological functions were differentially expressed varied for each species (Fig. 1B). For example, during growth *in planta*, adhesion– and motility-associated proteins increased in CPU and HRO, but decreased in SMA and ELU. Iron storage and acquisition-associated proteins similarly varied between species *in planta* in that they increased in BPI, HRO, PPU, and SMA and decreased in CIN and ELU. Interestingly, transport-associated proteins increased in all species when grown *in planta*.

**Figure 1.**
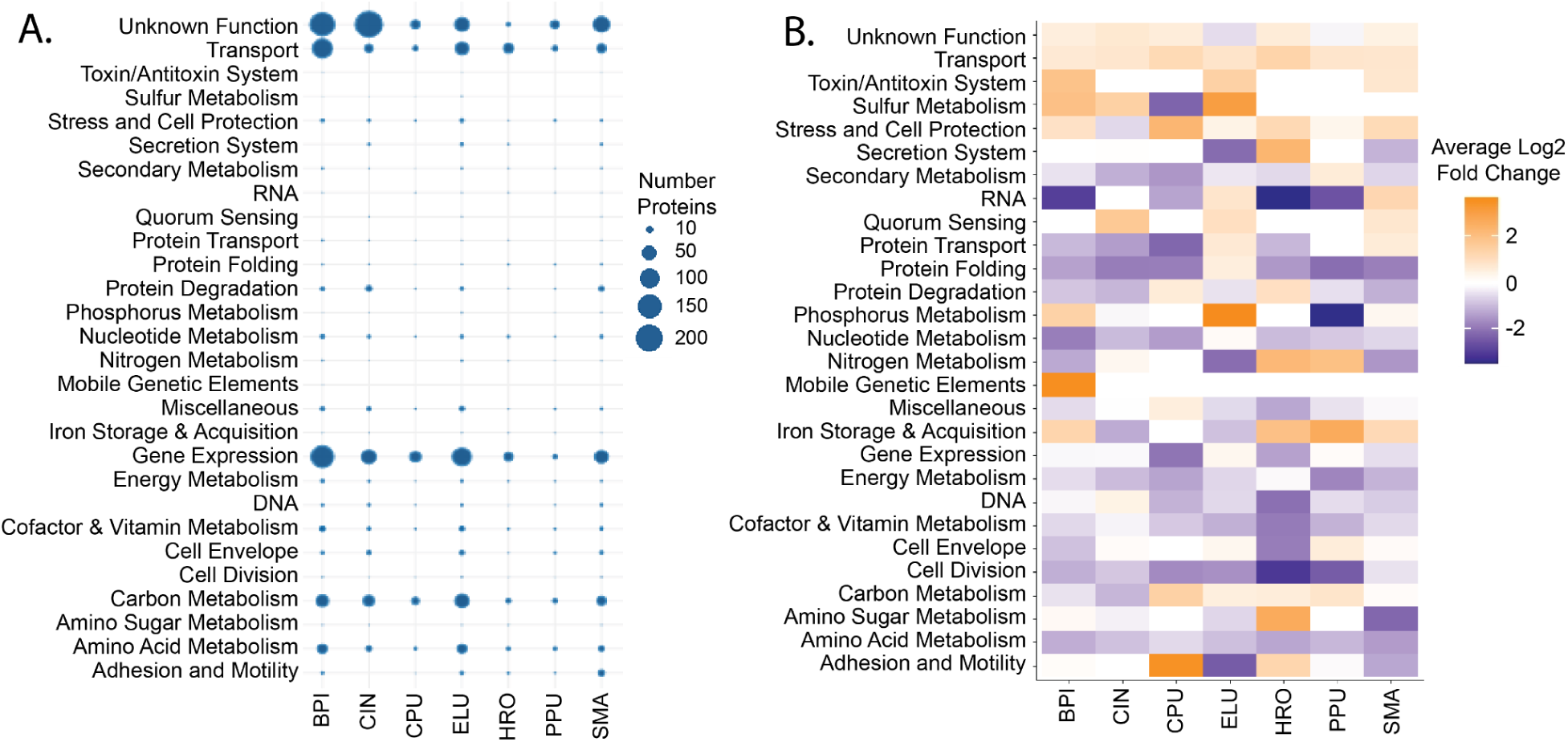
Protein abundances and differential abundance varied greatly between species when grown *in planta*. A) Number of proteins whose abundances significantly differed between *in planta* and *in vitro* growth. Broad functional categories are listed on the y-axis and individual species are listed on the x-axis (3-5 biological replicates) (Student’s T-Test, Benjamini-Hochberg adjusted p-value (*q*) < 0.05). B) Averages of log_2_ fold changes of all bacterial proteins whose abundances significantly differed between *in vitro* and *in planta* conditions (Student’s T-Test, Benjamini-Hochberg adjusted p-value (*q*) < 0.05), categorized by broad function. Fold change greater than 0 indicates a higher abundance *in planta*, and a fold change less than 0 indicates a higher abundance *in vitro*.

Although the broad functional categories provide an initial overview of functions that are most responsive to *in planta* growth, the detailed functional categories provide a clearer picture of the specific metabolism and physiology of each of the seven species *in planta* (Fig. 2). For example, within the “Carbon Metabolism” broad category, there are 29 detailed categories, which include pathways such as arabinose metabolism, inositol metabolism, and polysaccharide degradation. Here, it becomes evident that specific carbon metabolism pathways were more abundant *in planta* in each of the seven species (Fig. 2); this contrasts with the broad-level analysis of carbon metabolism-associated proteins where increased *in planta* abundances were detected only in CPU, ELU, HRO, and PPU (Fig. 1B). Similarly, in the broad-level analysis, HRO was the only species with increased *in planta* expression of secretion systems (Fig. 1B). However, when examining individual secretion systems, ELU and SMA showed increased expression of Type II Secretion System (T2SS) proteins (Fig. 2), while also showing decreased expression of T6SS proteins.

**Figure 2.**
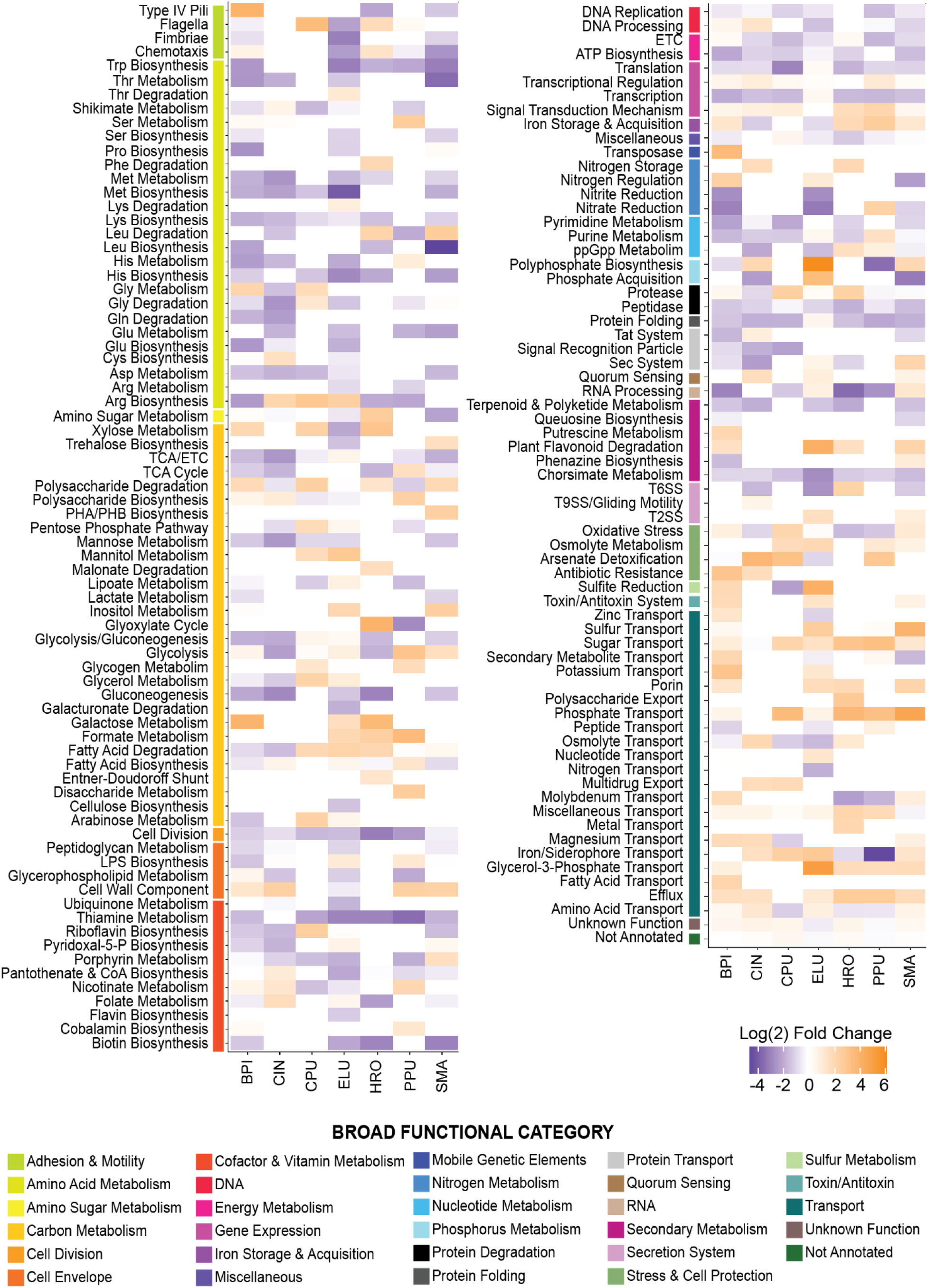
Differential expression of specific functional categories varied greatly between species in response to *in planta* growth. Averages of log_2_ fold changes of bacterial proteins whose abundances significantly differed between *in vitro* and *in planta* conditions (Student’s T-Test, Benjamini-Hochberg adjusted p-value (*q*) < 0.05), categorized by detailed and broad function. Fold change greater than 0 indicates a higher abundance *in planta*, and a fold change less than 0 indicates a higher abundance *in vitro*.

We therefore focused our analysis and follow-up work on detailed functional categories, with a particular focus on functions that were upregulated *in planta*. Below, we discuss results from proteins whose functions could potentially mediate (1) physical plant-bacterial interactions (Carbon Metabolism, Secretion Systems, Adhesion and Motility), (2) bacterial provisions to the plant (Phosphorus Metabolism), (3) general bacterial response systems (Transporters), and (4) proteins whose functions could be newly identified as plant-associated (Unknown Functions).

### Carbon metabolism pathways showed significant shifts in expression *in planta*

#### *Curtobacterium pusillum* (CPU) and *Herbaspirillum robiniae* (HRO) hemicellulose degradation enzymes were more abundant *in planta* and CPU was capable of using hemicellulose as a primary carbon source *in vitro*

We found that hemicellulose degradation enzymes increased in abundance in CPU and HRO *in planta* (Fig. 3A). Hemicelluloses are structural components of plant cell walls and include a range of polysaccharides, including xyloglucans, xylans, mannans, and glucomannans, among others (35). In CPU, α-L-arabinofuranosidase abundance was significantly higher *in planta* than *in vitro* (Fig. 3A). This enzyme cleaves arabinose sidechains from arabinoxylans, which are a common form of hemicellulose. Xylose isomerase, arabinose isomerase, ribulose-phosphate-3-epimerase, and ribose-5-phosphate isomerase (Supplementary Data 2) also had higher abundances *in planta* than *in vitro*, indicating that arabinose and xylose from arabinoxylan were further metabolized via the pentose phosphate pathway.

**Figure 3.**
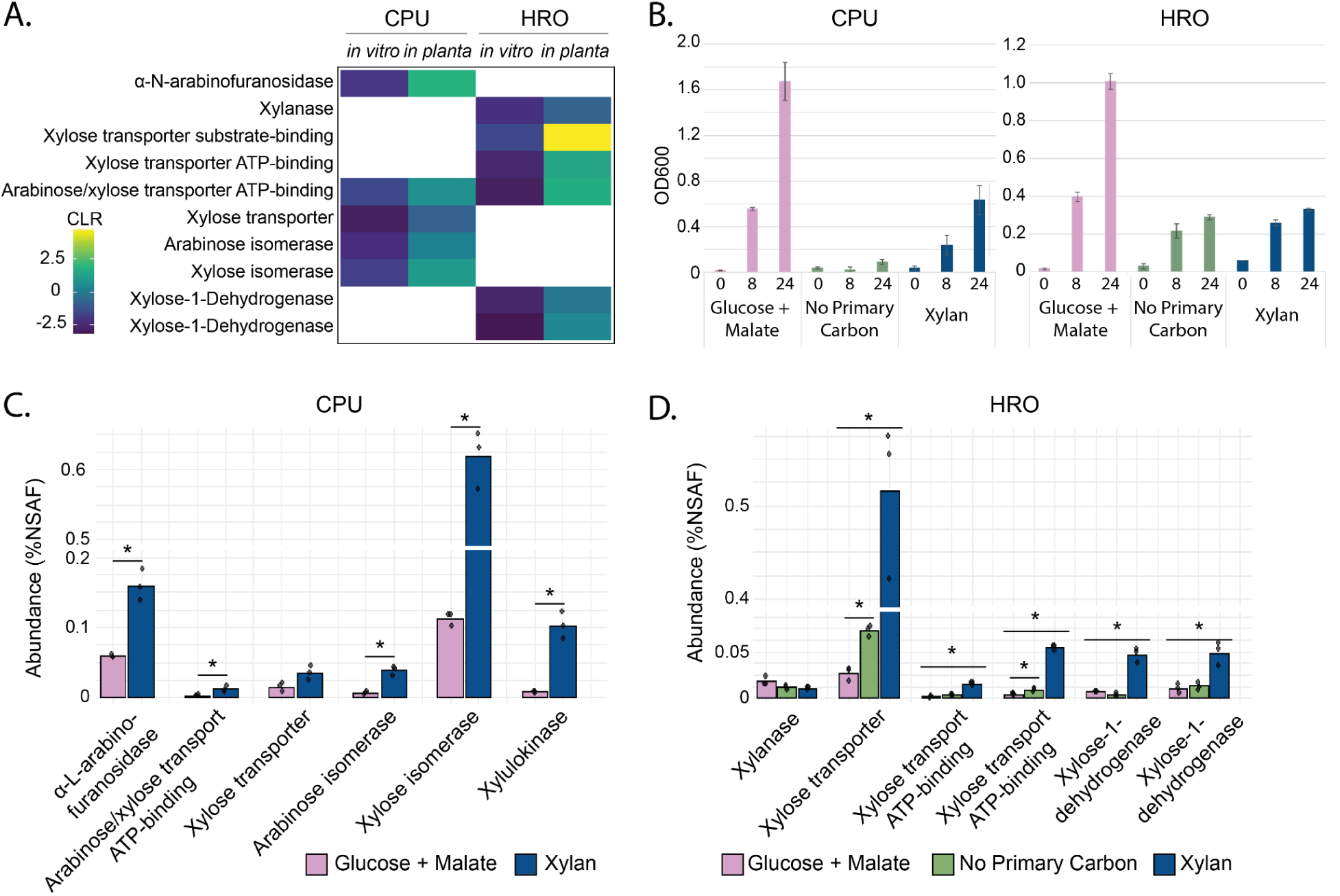
Hemicellulose-degrading enzymes in HRO and CPU were in higher abundance *in planta* and were expressed when grown on hemicellulose *in vitro*. A) Centered log-ratio transformed abundances of hemicellulose degradation-associated proteins that were significantly more abundant *in planta* in CPU and HRO (Student’s T-test, Benjamini-Hochberg adjusted p-value (*q*) < 0.05). White blocks indicate proteins that were either not identified or not significantly differentially abundant. B) OD_600_ values of CPU and HRO after 0, 8, and 24 hours of growth in minimal media with different primary carbon sources (n = 3). C) Average relative abundances (% normalized spectral abundance factor (NSAF)) of hemicellulose degradation-associated proteins when CPU was grown in minimal medium with glucose and malate or xylan (n = 3). D) Average relative abundances (%NSAF) of hemicellulose degradation-associated proteins when HRO was grown in minimal medium with glucose and malate, no primary carbon source, or xylan. The two xylose-1-dehydrogenases indicated are homologs. Asterisks indicate a Benjamini-Hochberg corrected p-value (*q*) < 0.05 (Student’s T-test). Error bars indicate standard deviation (n = 3).

In HRO, the abundance of a putative xylanase, which hydrolyzes xylan chains to xylose monomers, was significantly higher *in planta* than *in vitro*. Xylose transporter substrate– and ATP-binding proteins were additionally more abundant *in planta*, as well as xylose-1-dehydrogenase, which is involved in downstream xylose degradation. These indicate that HRO may also degrade hemicellulose.

To test whether CPU and HRO were capable of growing on hemicellulose, we performed an *in vitro* growth assay in which xylan was the primary carbon source. We grew CPU and HRO in three different versions of the minimal medium (MM): with glucose and malate, with xylan, and without any primary carbon source. Amino acids were also present in the medium (“secondary carbon source”; Supplemental Materials and Methods). We then used proteomics to determine whether the hemicellulose-degrading enzymes detected *in planta* were also present at increased levels when CPU and HRO were grown on xylan *in vitro*.

We found that CPU grew in the minimal medium with xylan and showed minimal growth when grown with no primary carbon source (Fig. 3B) (the growth that was seen without a primary carbon source was likely due to the amino acids present in the medium). Given the limited growth in the medium without a primary carbon source, we were unable to collect cells for proteomic analysis for this condition. However, we were able to compare abundances of hemicellulose proteins when CPU was grown with xylan and when it was grown with glucose and malate. We found increased abundances of five of the six putative hemicellulose catabolism proteins we had identified *in planta* when grown on xylan compared to when grown on glucose and malate (Fig. 3C). These proteins included ɑ-L-arabinofuranosidase, arabinose/xylose ABC transport ATP-binding protein, arabinose isomerase, xylose isomerase, and xylulokinase, further supporting the hypothesis that CPU degrades hemicellulose in the maize root environment. Altogether, our results showing that CPU hemicellulose degradation enzymes were more abundant *in planta*, that CPU was able to grow on hemicellulose as a primary carbon source *in vitro*, and that the same hemicellulose degradation enzymes were more abundant when grown on hemicellulose suggest that CPU does indeed degrade hemicellulose in the maize rhizosphere. However, the exact location of hemicellulose degradation is unclear. It has previously been reported that cell wall degradation can contribute to bacterial colonization of plant tissues (36, 37), which could suggest that CPU may colonize the endosphere. However, given that hemicelluloses are also available in the rhizosphere via sloughed off root cells (38, 39) and plant exudates (40, 41), CPU may also scavenge hemicellulose in the rhizosphere.

We additionally found that HRO grew equally well in the minimal medium with xylan as the primary carbon source and in the minimal medium without a primary carbon source (Fig. 3B), likely due to the presence of amino acids in the medium. It is therefore difficult to conclude whether HRO catabolizes xylan *in vitro*. However, when comparing proteins expressed in the presence of xylan to those expressed with no primary carbon source or with glucose and malate, we did find increased abundances of some of the hemicellulose catabolism enzymes that had been in higher abundance *in planta*, including xylose transporters and xylose-1-dehydrogenase (Fig. 3D). Notably, abundance of the putative xylanase did not change when xylan was present, indicating that it may have been mis-annotated and is not a true xylanase. Alternatively, it could be a true xylanase and its upregulation may not simply depend on the presence of xylan. Other factors such as the presence of amino acids in the medium may inhibit its regulation. Future work could test for xylanase activity using enzyme assays.

#### Fatty acids may be used as a carbon source by *Curtobacterium pusillum* (CPU) and *Herbaspirillum robiniae* (HRO) *in planta*

We found proteins from the fatty acid β-oxidation cycle that were significantly more abundant when CPU and HRO were grown *in planta* versus *in vitro* (Fig. 4A) (with the exception of acetyl-CoA C-acyltransferase, which, while not statistically significant, was also more abundant in both CPU and HRO *in planta*). This pathway is used in fatty acid catabolism, oxidizing fatty acids to acetyl-CoA, which can then be fed into the TCA cycle. Furthermore, the glyoxylate cycle, which funnels acetyl-CoA from β-oxidation into central carbon metabolism, was increased in HRO *in planta*: isocitrate lyase, which catalyzes the first step of the glyoxylate cycle, was significantly more abundant *in planta* than *in vitro*, and malate synthase, which catalyzes the second step, though not statistically significant, was also increased *in planta* (Fig. 4B) (42).

**Figure 4.**
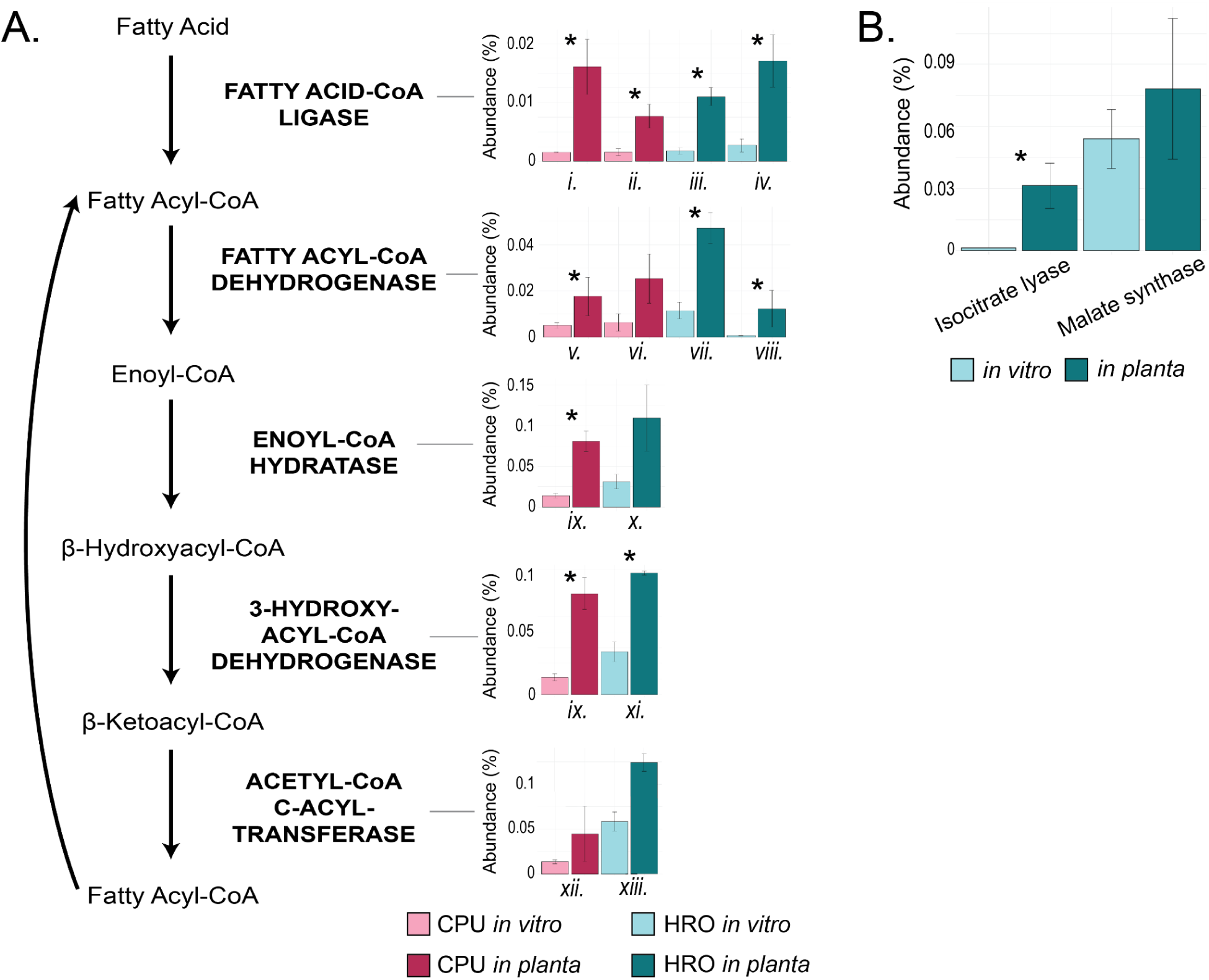
Fatty acid catabolism enzymes had higher abundances when *C. pusillum* and *H. robiniae* were grown *in planta* than *in vitro*. A) Abundances (% NSAF) of fatty acid β-oxidation cycle enzymes, with enzyme homologs indicated by Roman numerals. *i*) CPU_079236941.1, *ii*) CPU_079238824.1, *iii*) HRO_079214242.1, *iv*) HRO_079218280.1, *v*) CPU_079238788.1, *vi*) CPU_099051966.1, *vii*) HRO_079217199.1, *viii*) HRO_079218538.1, *ix*) CPU_079235882.1, *x*) HRO_079218255.1, *xi*) HRO_079217198.1, *xii*) HRO_079217197.1, *xiii*) CPU_079235883.1. Note that the enzymes listed as 3-hydroxyacyl-CoA dehydrogenase and enoyl-CoA hydratase for CPU (CPU_079235882.1) in Fig. 4A are the same enzyme; this enzyme showed homology to FadB, which contains two active sites: one 3-hydroxyacyl-CoA dehydrogenase domain and one enoyl-CoA hydratase domain, enabling it to perform both reactions (45). Error bars indicate standard deviation. B) Abundances (%NSAF) of glyoxylate cycle enzymes. Asterisks indicate a Benjamini-Hochberg corrected p-value (*q*) < 0.05 (Student’s T-test) when comparing CLR-transformed abundance values of proteins between *in vitro* and *in planta* conditions. Error bars indicate standard deviation.

Taken altogether, our data suggest that fatty acids are a potential carbon source for HRO and CPU *in planta*. Similar findings have been reported by Hemmerle *et al.* (12), who found an increase in *Rhizobium* β-oxidation cycle enzymes during co-inoculation with another bacterial species in the *Arabidopsis* phyllosphere, and by Van Dijk and Nelson (43), who suggested that competition for plant-derived fatty acids plays a role in biological control of *Pythium* by *Enterobacter* in cotton seedlings. Zhalnina *et al.* (44) additionally found that plant-derived fatty acids were depleted when bacteria were grown in *Avena* root exudates. However, there is much yet unknown about microbial consumption of fatty acids, including fatty acid type (e.g. length, saturation) and source (e.g. exudates, host cell membranes), particularly in the rhizosphere. This highlights a gap in our understanding of microbial physiology *in planta*, as it is possible that fatty acids are an important carbon source for plant-associated bacteria.

### Phosphate solubilization enzyme abundance *in planta* varied between species

When ELU was grown on the maize root, we found that phosphorus metabolism-related proteins were more abundant than when grown *in vitro* (Fig. 1B) In particular, we observed a higher abundance of alkaline phosphatase *in planta* (Fig. 5A). Alkaline phosphatases are involved in liberating phosphate from organic compounds and polyphosphates, allowing organisms to scavenge phosphate from the environment when free phosphate is limited (46). The higher abundance of this enzyme in ELU *in planta* could therefore suggest a phosphate limitation in the root environment. Interestingly, however, we observed the opposite trend in CIN, CPU, and SMA wherein their alkaline phosphatases were actually less abundant *in planta* than *in vitro*. Since all plants were watered with the same amount and concentration of Murashige-Skoog medium, these contrasting patterns point to species-specific regulation of phosphate solubilization. Three of the four alkaline phosphatases (encoded by CIN, ELU, and SMA) contained predicted signal peptides and cleavage sites (SignalP (47)), indicating that they are secreted and likely function extracellularly.

**Figure 5.**
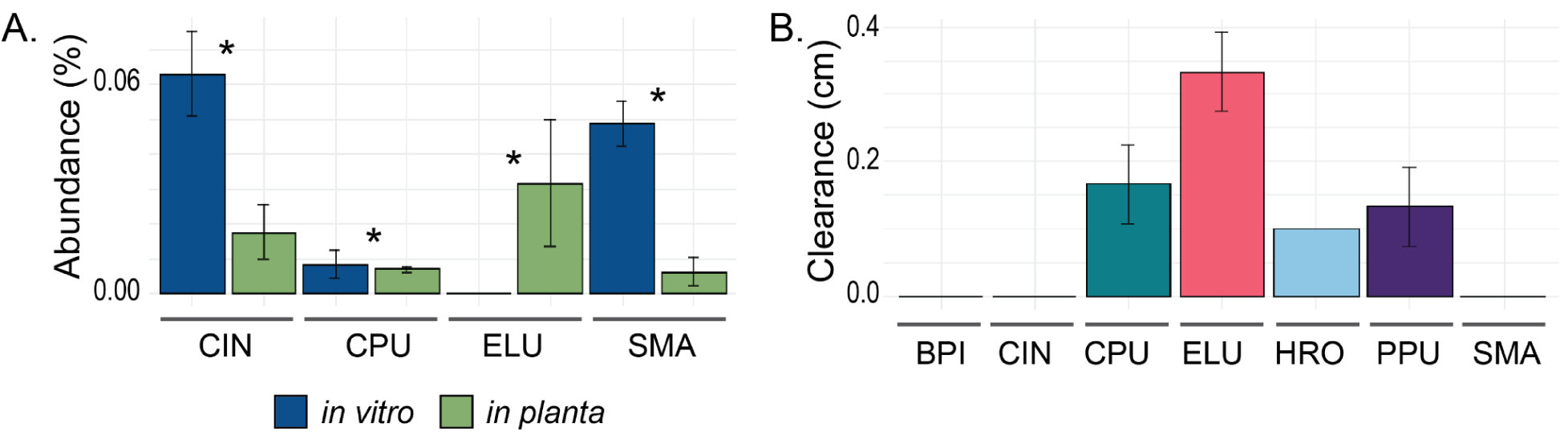
*Enterobacter ludwigii* (ELU) encoded alkaline phosphatase was more abundant *in planta* and ELU is capable of phosphate solubilization *in vitro*. A) Relative abundance (% NSAF) of significantly differentially abundant alkaline phosphatases in ELU, CIN, and SMA when grown *in vitro* and *in planta* (Student’s T-Test; Benjamini-Hochberg corrected p-value (*q*) < 0.05). Error bars indicate standard deviation. B) Zone of clearance (cm), indicative of phosphate solubilization, when each species was grown on Pikovskaya agar after 5 days. Error bars indicate standard deviation.

We confirmed that ELU and CPU are also capable of phosphate solubilization *in vitro* on Pikovskaya agar (Fig. 5B), supporting previous studies that have shown that *Enterobacter* and *Curtobacterium* species solubilize phosphate (48, 49). This indicates that these alkaline phosphatases may contribute to phosphate solubilization *in planta*, increasing its availability for uptake by the maize root.

We also measured phosphate solubilization capability for the other five species on Pikovskaya agar (Fig. 5B), and surprisingly found that CIN and SMA did not solubilize phosphate in this assay. However, it has previously been shown that some bacteria are capable of phosphate solubilization in liquid media, but not agar (50), which may explain this observation. We additionally found that HRO and PPU solubilized phosphate, though we did not detect alkaline phosphatases in either proteome. We found that both species encode putative alkaline phosphatases in their genomes (WP_079230605.1 and WP_079226921.1 in PPU and WP_079216850.1 in HRO); while these may be mis-annotated, this could indicate that either these alkaline phosphatases were not expressed under our *in vitro* and *in planta* conditions, or that they were not expressed to levels high enough for detection. These two species may additionally solubilize phosphate via a separate mechanism, such as organic acid exudation (51). We found no phosphate solubilization on Pikovskaya agar by BPI, nor did we find any alkaline phosphatases encoded in its genome.

### Bacterial secretion systems were differentially regulated *in planta*

#### Both Type VI Secretion Systems in *Herbaspirillum robiniae* (HRO) are involved in colonization and growth on maize roots

T6SSs are protein secretion systems commonly found in Gram-negative bacteria that have been shown to play important roles in mediating host-microbe and microbe-microbe interactions (52–57). In plant-associated bacteria, the T6SS is found in pathogenic and beneficial taxa and can play roles in plant disease, plant colonization, and interactions with other microbes in the environment (55–58). Of the seven bacterial species investigated in this study, five encode genes for T6SSs, with PPU, CIN, and SMA each encoding one T6SS, and ELU and HRO each encoding two T6SSs. While most of the differentially abundant T6SS proteins in CIN, ELU, PPU, and SMA were found in lower abundances *in planta* compared to *in vitro*, proteins found in both HRO T6SS gene clusters were significantly more abundant *in planta* (Fig. 6A, 6B). This suggests that the T6SSs may play an important role in interactions between HRO and the maize root.

**Figure 6.**
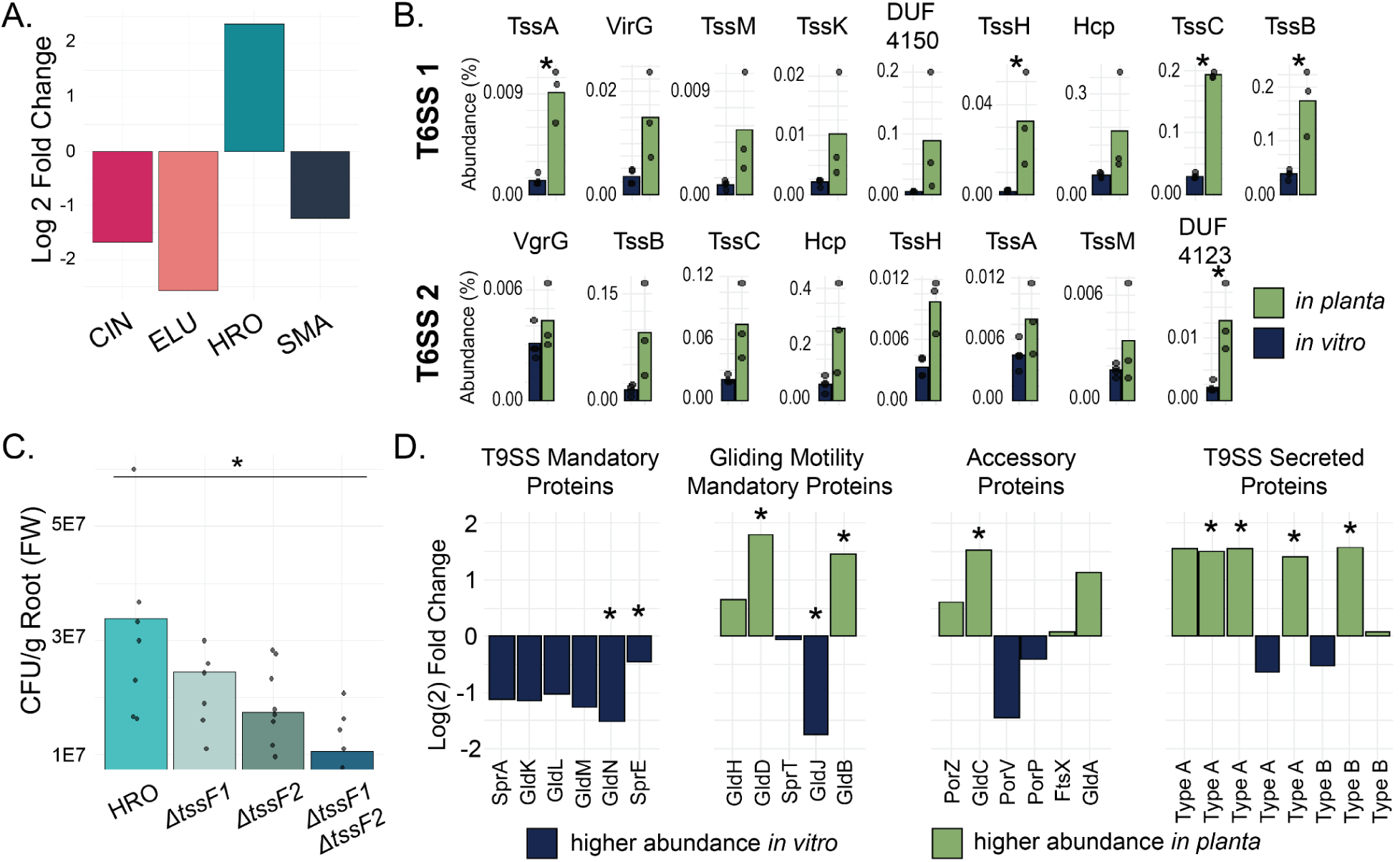
Secretion system expression during *in planta* growth. A) Average log(2) fold change (calculated with centered-log ratio transformed abundances) of differentially abundant T6SS proteins in each species for which T6SS proteins were detected (n = 3-5). A log(2) fold change greater than 0 indicates a higher abundance *in planta*, while a log(2) fold change less than 0 indicates a higher abundance *in vitro*. B) Abundances (% NSAF) of detected HRO T6SS proteins (n = 3-4). Asterisks indicate a Benjamini-Hochberg corrected p-value (*q*) < 0.05 (Student’s T-test) when comparing CLR-transformed abundance values of proteins between *in vitro* and *in planta* conditions. C) Mean colony forming units (CFU) per gram of root fresh weight (FW) of HRO T6SS mutants when grown on maize roots for 14 days (n = 8-9). Asterisk indicates a p-value < 0.05 (Student’s T-test) when comparing each knockout mutant to the wildtype (HRO). D) Log(2) fold change (calculated with centered-log ratio transformed abundance) of differentially abundant T9SS and gliding motility proteins in CIN. A log(2) fold change greater than 0 indicates a higher abundance *in planta*, while a log(2) fold change less than 0 indicates a higher abundance *in vitro*. Asterisks indicate a Benjamini-Hochberg corrected p-value (*q*) < 0.05 (Student’s T-test) when comparing CLR-transformed abundance values of proteins between *in vitro* and *in planta* conditions.

Notably, we did not identify all proteins classically associated with T6SSs. The T6SS is a multi-protein complex where certain proteins are present in high copy numbers (e.g. Hcp, TssB, TssC), while others exist in only one or few copies (e.g. TssF, TssK), making the latter group of proteins more difficult to detect. Furthermore, some T6SS proteins are fully membrane-bound, while others are secreted or intracellular, and these membrane-bound proteins can be harder to detect than soluble proteins.

In order to investigate the *in planta* role of the HRO T6SSs, we generated three T6SS knockout mutants by deleting the *tssF* gene in each (single knockouts) and both (double knockout) of the gene clusters, as deletion of *tssF* has previously been shown to reduce T6SS function (52, 59). This resulted in the following mutants: *ΔtssF1*, *ΔtssF2*, and *ΔtssF1ΔtssF2*. We inoculated each of these mutants and the wildtype strain individually on sterile maize seedlings and grew the plants for two weeks to determine whether functional loss of the T6SS would result in reduction in colonization. While we saw some (non-significant) reduction in colonization by the single *tssF* mutants compared to the wildtype, the strongest effect was seen in the double mutant, *ΔtssF1ΔtssF2*, with an approximately 3-fold reduction in abundance (*p* < 0.05) (Fig. 6C). This indicates that the use of both T6SSs has a positive effect on colonization and growth of HRO on the maize root.

To determine whether this effect was due to a general growth defect in HRO when *tssF1* and *tssF2* were knocked out, we grew each of the *tssF* mutants and the wildtype in tryptic soy broth (TSB) and the minimal medium for 24 hours (Fig. S1, Fig. S2). While we found no significant changes in growth between the knockout mutants and the wildtype in TSB, the double mutant, *ΔtssF1ΔtssF2*, grew to a significantly higher OD_600_ than the wildtype in the minimal medium (Student’s T-Test; p < 0.05). This suggests that expressing both T6SSs comes at a fitness cost to HRO under certain conditions. This contrasts with the *in planta* results, indicating that the effects seen *in planta* were not due to a general growth defect when either or both *tssF* copies were deleted.

The increased abundance of T6SS proteins in HRO *in planta*, along with the reduced colonization by the T6SS mutants, suggests that these secretion systems contribute to HRO colonization and persistence on maize roots. This is consistent with previous studies that have shown that T6SSs contribute to colonization and persistence by both pathogenic and non-pathogenic plant-associated bacteria (60, 61). In the context of direct plant-microbe interactions, T6SSs have been implicated in diverse *in planta* functions, including biofilm formation, siderophore production, and metal acquisition (60, 62, 63). T6SSs are also known to be involved in interbacterial competition and killing (55, 56), although this role is less relevant in this study, as each species was inoculated on sterile seedlings individually. Further work is needed to identify the specific proteins secreted by the HRO T6SSs *in planta* and to understand how they influence root colonization and persistence and, potentially, interactions with other microbes under more complex conditions.

#### Type IX Secretion System proteins are found to be expressed at different levels *in planta* in *Chryseobacterium indologenes* (CIN)

The T9SS is involved in secretion of a large diversity of proteins, including those involved in polysaccharide degradation, motility, and adhesion, among others (64, 65). While the role of the T9SS in plant-associated microbes is still largely unknown, there is evidence that it may play a role in colonization of seeds and plant roots (66). Using T9GPred, NCBI PGAP, and UniProt, we identified 29 genes encoding proteins essential for the T9SS and gliding motility, as well as accessory and secreted proteins, in CIN.

We detected 25 of these proteins in the CIN proteomes and found that 10 were significantly differentially abundant either *in vitro* or *in planta* (Fig. 6D). Interestingly, all T9SS-mandatory proteins were found in lower abundance *in planta* than *in vitro* (Fig. 6D), while four of the eight predicted T9SS-secreted proteins were significantly more abundant *in planta*. Previous studies have shown that T9SS substrates accumulate intracellularly or within the periplasm when T9SS components are knocked out, suggesting that T9SS substrate gene expression is not dependent on T9SS component expression (67–69), which could explain the pattern seen in this study. We used InterProScan to identify domains that may indicate function of these potentially secreted proteins, but were unsuccessful. Taken altogether, these findings indicate that the predicted T9SS-secreted proteins may play a role independent of the T9SS machinery and/or may be secreted through an alternative pathway, leaving the role of the T9SS in CIN growth on maize roots unclear.

We additionally found that gliding motility-mandatory proteins and accessory proteins were differentially abundant, though condition-dependent (Fig. 6D). Based on the contrasting results, we are unable to speculate as to its role in interactions with the maize root.

### Transporter proteins were more abundant in all seven bacterial species when grown *in planta*

Transporter proteins were significantly more abundant *in planta* than *in vitro* across all seven species studied (Fig. 1B, Fig. 2). The expression patterns of these transporters can provide useful insight into the physiologies and substrate preferences of bacteria, as well as nutrient availability in a given environment. In total, we annotated 570 differentially abundant transporter proteins across all seven species and categorized them into 21 different functional categories (Supplementary Data 2). Among these, we found that three transporter types were particularly abundant, each comprising more than 1% of the proteome in at least one species: amino acid transporters, porins, and sugar transporters (Fig. 7A). Miscellaneous transporters also composed a sizable portion of the proteome in BPI, CIN, and SMA. However, the proteins that were annotated as belonging to “Miscellaneous Transport” were largely transporter components for which we were unable to predict a substrate, and thus, we do not discuss these in depth.

**Figure 7.**
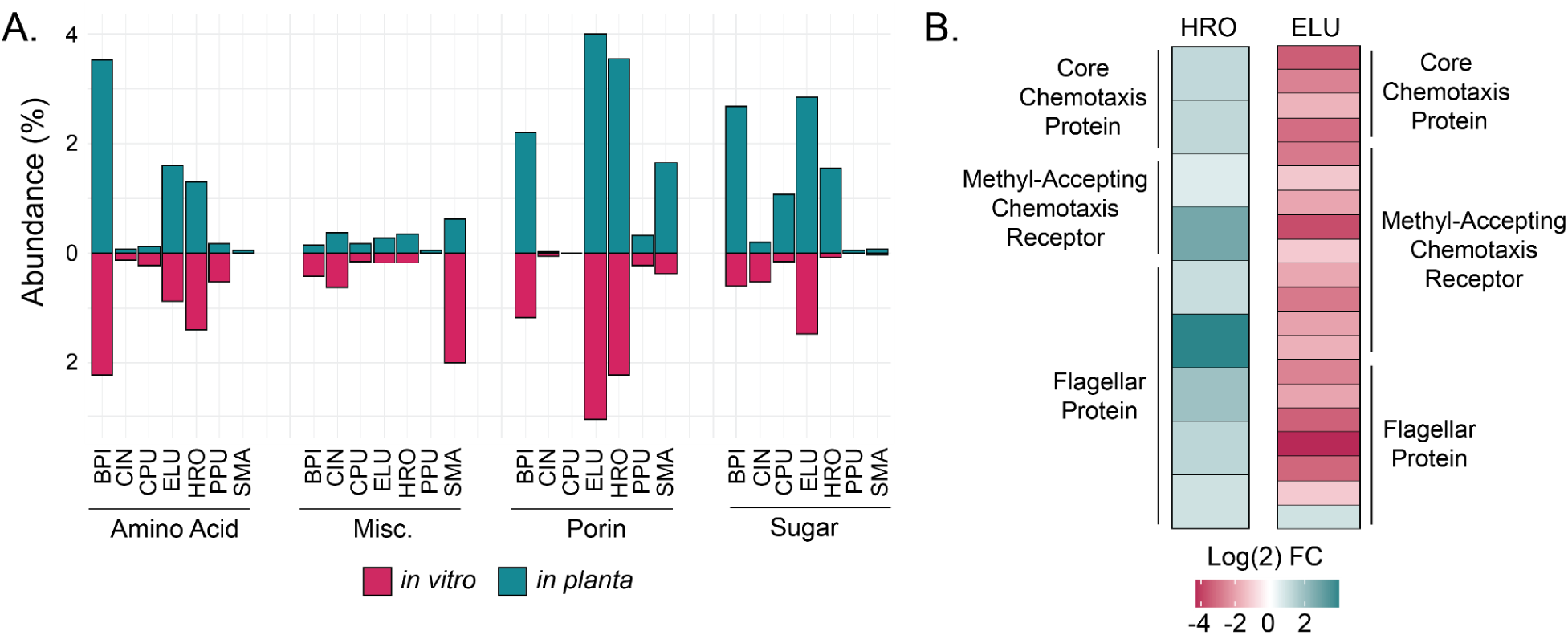
Proteins from four transporter categories composed at least 1% of the proteome in at least one of the bacterial species studied. Chemotaxis and motility proteins were differentially abundant in *H. robiniae* and *E. ludwigii*. A) Summed abundance (% NSAF) of significantly (Student’s T-Test; Benjamini-Hochberg corrected p-value (*q*) < 0.05) differentially abundant transporter proteins across all seven bacterial species. B) Log(2) fold change (of CLR-transformed abundance values) of significantly differentially abundant chemotaxis and flagellar proteins in *H. robiniae* and *E. ludwigii*. A log(2) fold change greater than 0 indicates a higher abundance *in planta*, while a log(2) fold change less than 0 indicates a higher abundance *in vitro* (Student’s T-Test; Benjamini-Hochberg corrected p-value (*q*) < 0.05).

#### Amino Acid

Abundances of amino acid transporter proteins varied across each of the seven species: differentially expressed amino acid transporter proteins in BPI, ELU, and HRO were particularly abundant and high in number, while CIN, CPU, PPU, and SMA had markedly less abundant transporter proteins (Fig. 7A).

In ELU, high-affinity arginine, branched-chain amino acid, cystine, glutamine, and histidine transporter proteins (both ATP-binding and substrate-binding) were significantly more abundant *in planta* than *in vitro* (Supplementary Data 2). This effect could be due to the fact that the minimal medium used in the *in vitro* condition included arginine, leucine, isoleucine, cystine, and glutamine at concentrations of 200-350 µmol/L, while it is estimated that the concentrations of individual amino acids in the rhizosphere are approximately 100-1,000x lower (70).

Surprisingly, it was difficult to make observations regarding amino acid uptake in HRO and BPI in the root environment, as transporter components were not consistently differentially abundant in the same condition (Supplementary Data 2). For example, among BPI branched-chain amino acid ABC transporter proteins, two substrate-binding proteins and one permease were more abundant *in planta*, while two ATP-binding proteins were more abundant *in vitro*. These results may be confounded by the fact that it can be difficult to assign transporter substrates by the ATP-binding subunit, as the ATP-binding subunit does not bind directly to the transporter substrate. Thus, it remains unclear which amino acids HRO and BPI may be scavenging in the plant environment.

#### Porins

Porins were significantly more abundant *in planta* across multiple species, particularly BPI, ELU, HRO, and SMA, suggesting an important role in bacterial survival in the maize rhizosphere. Porins are outer membrane proteins that generally facilitate passive diffusion of small molecules across the cell membrane, although it has been suggested that they can be cation– or anion-specific under certain conditions (71). In plant-associated bacteria, porins have been associated with nutrient uptake, biofilm formation, antibiotic sensitivity, and stress resistance (71–73). While similarly high abundances of bacterial porins have been found in the phyllosphere and rhizosphere (13, 74, 75), the lack of porin specificity and broad range of potential functions currently makes it difficult to draw conclusions as to the role of porins in BPI, ELU, HRO, and SMA *in planta*.

#### Sugar

Of the 82 differentially abundant sugar transport-related proteins, 69 were more abundant *in planta*. These proteins belonged to a variety of transporter families and/or mechanisms, including MFS, ABC, Sus, and PTS transport mechanisms. While we were unable to identify potential substrates for all transporters, we did find a number of 5-carbon, 6-carbon, disaccharide, and polysaccharide transporters, indicating that these species consume a large diversity of substrates.

Notably, all differentially abundant sugar transporters identified in CIN belong to the starch utilization system (Sus), which is widespread in the phylum Bacteroidetes. We identified three SusD homologs upregulated *in vitro* and four upregulated *in planta*, as well as one SusE/F homolog upregulated *in vitro* (Supplementary Data 2). SusDEF proteins are outer membrane-bound starch-binding proteins that have been shown to be heavily involved in a cell’s ability to grow on different starches (76). We additionally identified two SusC homologs upregulated *in vitro* and four upregulated *in planta*. These are TonB-dependent transporters responsible for the translocation of maltooligosaccharides into the periplasm (76).

These results suggest that sugar uptake is a key component of bacterial adaptation to the maize root environment, with multiple species exhibiting increased expression of transporters for a wide range of carbon sources. The differential expression of SusCDEF components in CIN further supports the idea that root-associated bacteria dynamically regulate starch utilization in response to environmental conditions.

### Chemotaxis and motility protein abundances were highly species-specific *in* planta

Two distinct patterns of chemotaxis and motility gene expression were present between HRO and ELU. In HRO, all significantly differentially abundant chemotaxis and flagellar proteins were more abundant *in planta* (Fig. 7B), suggesting that HRO increases motility in response to specific environmental cues in the root environment. In contrast, ELU showed the opposite pattern: all thirteen differentially abundant chemotaxis proteins were significantly more abundant *in vitro*, as were six of the seven differentially abundant flagellar proteins (Fig. 7B). This indicates a shift in ELU from a motile lifestyle *in vitro* to a less motile lifestyle *in planta*.

While many of the classic chemotaxis signaling cascade pathway proteins were present in HRO and ELU, such as CheA, CheV, CheW, and CheZ, there were also a number of methyl-accepting chemotaxis proteins and receptors whose ligands are unknown. As such, it is difficult to predict the chemoattractants to which these microbes may respond.

Chemotaxis and motility have often been associated with increased plant colonization capability of bacteria (77–79). However, it has been suggested that some bacteria require chemotaxis and motility functions specifically during early colonization, but form biofilms upon colonization (78). Given that the plants in this experiment were grown for two weeks, we would not have detected functions relevant to early colonization, but rather those involved in continued persistence in the root environment. It is therefore possible that chemotaxis and motility are still involved in ELU colonization of maize roots, but not highly expressed once colonization is established. While we did not identify differentially abundant biofilm formation-related proteins in ELU, it is possible that ELU forms a biofilm *in planta*, given that other *Enterobacter* species have been shown to form biofilms (80, 81) in different environments.

Thus, while chemotaxis and motility are commonly associated with enhanced plant-microbe interactions, the contrasting patterns of chemotaxis and motility gene expression between HRO and ELU suggest that their roles in root-associated bacteria are highly species-specific and potentially context-dependent.

### Cell division protein abundance varied greatly by protein, suggesting heterogeneity in growth phase dynamics within populations

We found that cell division proteins were, on average, more abundant *in vitro* than *in planta* (Fig. 1B). Given that cells from the *in vitro* condition were harvested at mid-log phase, this pattern could suggest that cell division occurred at a higher rate *in vitro* and that cells *in planta* may either be growing more slowly or have entered stationary phase. However, upon deeper analysis of each of the differentially abundant cell division proteins, the pattern is more complex (Fig. 8). For each of the seven species, some cell division proteins were more abundant *in planta* while others were more abundant *in vitro*, making it difficult to draw conclusions as to growth phase *in planta*. One possible explanation is that bacterial populations along the primary root are not homogeneous; because cells were harvested from the entire primary root, the sampled population comprised bacteria residing in distinct microenvironments (82). As a result, some cells may be actively dividing, while others grow more slowly or enter stationary phase, depending on the location on the root.

**Figure 8.**
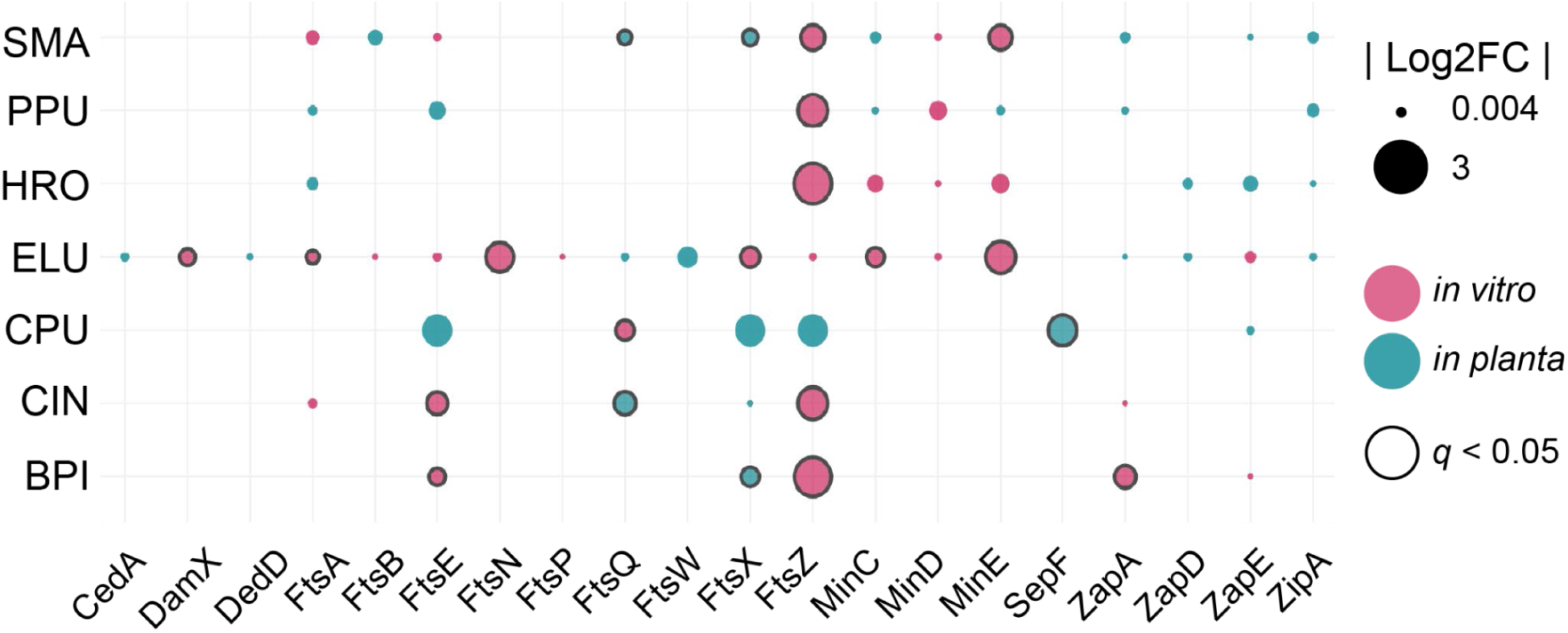
Log(2) fold change of cell division proteins. Size corresponds to log2FC. Pink indicates proteins that were more abundant *in vitro*, green indicates proteins that were more abundant *in planta*. Black outline indicates statistical significance (Student’s T-Test; Benjamini-Hochberg corrected p-value (*q*) < 0.05).

Previous experimental work has shown that cell counts of CPU and ELU increase by several orders of magnitude on the root over the course of 10-14 days, suggesting active growth during that time period by these two species (24, 26). In both studies, PPU counts decrease over these time periods, suggesting death, while the other four species show varying degrees of growth or decline between the two studies. This highlights an avenue for future work, as, to the best of our knowledge, the growth dynamics of bacteria in plant roots remain largely unknown, though some work has been done in the phyllosphere of *Arabidopsis* (83).

### Proteins of unknown function made up a large proportion of the differentially expressed proteins in each of the seven species

It is important to note that despite extensive manual and automated curation efforts, 2,192 of the 5,265 differentially abundant proteins could not be confidently annotated and/or categorized. These proteins were either assigned “Unknown Function” or were not annotated at all. This indicates that there remain a large number of microbial functions that may be highly relevant for growth and colonization of maize roots that are yet undiscovered. This highlights that abundant and strongly differentially abundant proteins with unknown functions should be prioritized as targets for biochemical and molecular functional characterization, as these may drive novel phenotypes.

## Conclusions

In this study, we used differential proteomics to investigate how seven maize root-associated bacterial species alter their gene expression when colonizing and persisting on the maize root. While the differential proteomic approach used in this study has uncovered many microbial functions related to growth in the maize rhizosphere, we recognize that there are limitations to our approach. Firstly, we studied plant-microbe interactions with mono-colonization experiments; in nature, microbe-plant interactions occur in complex environments with other organisms at play, and it has become increasingly clear that microbe-microbe interactions can have significant impacts on bacterial gene expression (12, 56, 84, 85). Based on this, we expect that microbe-microbe interactions will have strong impacts on microbial *in planta* gene expression, and such gene expression responses are not captured in our study. Secondly, while we carefully curated our protein annotations by using multiple databases and tools, we recognize that homology-based protein annotations are not always correct (86, 87). We successfully substantiated several functions that were differentially abundant by using *in vitro* assays, proteomics, and knockout mutants. While the task of confirming all differentially abundant functions is much too large for a single study, our results indicate that proteomic changes are a good indicator of changes in microbial functions. Thirdly, we were unable to confidently annotate a sizable portion of the proteins we identified. Thus, it seems likely that many functions related to maize colonization and growth remain uncharacterized in these species. Indeed, it has been suggested that 40-60% of genes cannot be confidently assigned functions (88), which has prompted work in recent years to develop high-throughput computational and experimental methods to assign functions to unknown proteins (88–90). Plant-associated bacteria are not exempt from this problem that faces all microbiologists, and must be included in future annotation efforts. Finally, we acknowledge that some observed differences in gene expression may be due to growth in a liquid medium versus a solid medium (maize root in a clay-sand mixture). However, we did design the minimal medium used in the *in vitro* condition to simulate the nutrient conditions of the root environment by using Murashige-Skoog as the base medium (this was also used to water the plants in the *in planta* treatments) and by using carbon sources readily found in maize root exudates (91). Moreover, several of the responses we observed – such as those related to plant polysaccharide degradation, phosphate solubilization, and T6SSs – are consistent with known mechanisms of plant-microbe interactions, and were supported experimentally. Thus, while we cannot fully exclude environmental effects unrelated to the plant, we are confident that many of the observed proteomic shifts reflect responses to growth on the maize root.

By investigating gene expression in single bacterial species grown *in vitro* and *in planta*, we were able to identify specific physiological and metabolic traits that enable colonization and growth in the maize root environment. These results provide several avenues for future research into microbe-host, microbe-microbe, and microbiota-host interactions. Our findings on the role of HRO T6SSs in rhizosphere growth raise further questions regarding the function of each T6SS and how they may interact with one another and with the plant host to increase colonization. Furthermore, given that T6SSs have frequently been associated with microbe-microbe interactions in various plant systems (55, 56, 92), understanding whether and how the T6SSs in HRO play a role in microbe-microbe interactions in the maize root environment would provide insights into the dynamics of microbial interactions within microbiota in the plant host context. Along these lines, as mentioned above, it is important to remember that host-microbe interaction studies must be scaled up in complexity in order to draw meaningful conclusions about host-microbe interactions *in situ*. Given that each of these bacteria belong to the same SynCom, we suggest that the next step would be to repeat a similar experiment in which bacterial functions are compared between mono-inoculation and co-inoculation with the entire SynCom to begin understanding how the dynamics and interactions within microbiota may impact and/or alter microbe-host interactions. Ultimately, it will be essential to study such plant-microbe interactions in increasingly complex communities to better understand the dynamics of interactions between different members *in situ*.

## Methods

### Culturing and inoculation of seven bacterial species for *in vitro* and *in planta* characterization

We used the isolates *Stenotrophomonas maltophilia* AA1 (SMA; DSM 114483), *Brucella pituitosa* AA2 (BPI; DSM 114565), *Curtobacterium pusillum* AA3 (CPU; DSM 114566), *Enterobacter ludwigii* AA4 (ELU; previously *E. cloacae*; DSM 114484), *Chryseobacterium indologenes* AA5 (CIN; DSM 114485), *Herbaspirillum robiniae* AA6 (HRO; DSM 114508), and *Pseudomonas putida* AA7 (PPU; DSM 114486) published by Niu *et al.* (24). We cultured all isolates in preparation for this experiment as described by Salvato *et al.* (27). Detailed methods can be found in SI Materials and Methods.

For the *in vitro* characterization portion of the experiment, we inoculated 1 mL of each washed overnight culture into 50 mL of a defined, minimal medium. We designed this minimal medium to consist of a base of 0.5x Murashige-Skoog (MS; Research Products International) augmented with glucose, malate, amino acids, and vitamins in order to grow the seven species in an environment as similar as possible to the *in planta* environment (SI Materials and Methods; Tables S2 and S3). We used four replicates (n = 4) for each species. We incubated cultures at 30°C in a shaker at 180 rpm; OD600 was measured immediately after 0, 2, 3, 4, 5, and 6 hours. When the cultures were at approximately mid-log phase, 2 mL of each culture was centrifuged at 8000 xg for 8 minutes, and each pellet was frozen at –80°C until protein extraction.

For the *in planta* characterization portion of the experiment, we diluted the remainder of each washed and re-suspended overnight culture from above to 10^7^ cells/mL, which was determined by OD600 using an OD-to-CFU standard curve (described in Salvato *et al.* (27)). 10 mL of the diluted cell suspension was added to 1 L of 0.5x Murashige-Skoog (final concentration of 10^5^ cells/mL). We sterilized *Zea mays* cv. Sugar Bun seeds (untreated; Johnny’s Selected Seeds) following a protocol described by Wagner *et al.* and Parnell *et al.* (23, 93). Sterile seeds were deposited individually in sterile WhirlPak bags (Nasco; SKU B01450) containing 50 mL of a sterile 1:1 clay (Pro’s Choice Rapid Dry, OIL-DRI)/sand (Multi-Purpose sand, Sakrete) mixture. Additional 50 mL of sterile 1:1 clay sand mixture were added on top of the seed. Seeds were watered with 65 mL (6.5 x 10^6^ cells) of the cell suspensions prepared above. Bags were sealed with AeraSeal (Millipore Sigma), and placed in a growth chamber at constant 25°C, with a cycle of 16 hours of fluorescent light and 8 hours of darkness for 14 days. For HRO, SMA, PPU, CIN, and CPU, we inoculated 6 bags, each containing one seed, to account for any seeds that did not grow or germinate, with the goal of obtaining 4-5 biological replicates. For BPI and ELU, we were not able to extract a sufficient amount of bacterial protein from a single plant, so we pooled two plants for each replicate. We therefore planted 20 bags, again to account for any seeds that did not grow or germinate, with the goal of having 10 plants that could be pooled for 5 replicates.

### Collection of bacterial cells from maize roots

After 14 days, plants were gently pulled from the WhirlPak bags. Roots were gently rinsed with dH_2_O to remove adhering clay/sand and the primary roots were cut from plants, weighed, and cut roughly into 1 cm pieces. We harvested bacteria from maize roots following the protocol described in Salvato *et al.* (27). Root fragments were placed in 1 mL sterile PBS with six sterile 3 mm glass beads and vortexed 3 times for 1 minute, with 10 seconds dwell time between each vortex. The resulting slurry was transferred to new centrifuge tubes and centrifuged at 15,000 xg for 7 minutes. Supernatant was removed and pellets frozen at –80°C.

### Protein extraction and peptide preparation

Bacterial pellets were removed from –80°C and thawed at room temperature. SDT lysis buffer (4% [wt/vol] SDS, 100 mM Tris-HCl, pH 7.6, 0.1 M DTT) was added to the pellets at a ∼10:1 ratio (e.g. 500 µL SDT for a 50 µL pellet). Suspensions were heated for 5-10 minutes at 95°C in a heat block. The Gram-positive species, CPU, was sonicated prior to heating in SDT using a Qsonica Q700 with three cycles of 30 seconds at 10% amplitude on and 1 minute off in order to disrupt the cell wall. Samples were then centrifuged for 5 minutes at 21,000 xg to remove cell debris.

We prepared peptides from the resulting protein extracts for CIN, CPU, HRO, SMA, and PPU using a filter-aided sample preparation (FASP) protocol adapted from Wisniewski *et al*. (94, 95). Detailed methods can be found in SI Materials and Methods.

We had difficulty attaining sufficiently high quality chromatograms for ELU and BPI using the FASP protocol and thus used a suspension trapping (S-Trap) sample preparation protocol with a few modifications to prepare peptides from these species (96). Details can be found in SI Materials and Methods. We used the Pierce microBCA assay (Thermo Fisher Scientific) to determine peptide concentrations.

### LC-MS/MS

HRO, SMA, PPU, CIN, and CPU samples were analyzed by one-dimensional LC-MS/MS as described by Mordant and Kleiner, and samples were blocked and randomized according to the method published by Oberg and Vitek (95, 97). Details can be found in SI Materials and Methods.

ELU and BPI samples were analyzed using a similar method to that above, but with a few modifications. These are described in SI Materials and Methods. The mass spectrometry proteomics data has been deposited to the ProteomeXchange Consortium via the PRIDE (98) partner repository with the dataset identifier PXD064252.

### Protein databases

We constructed protein sequence databases for protein identification by combining the protein sequences from each bacterial species with the *Zea mays* protein sequences (we used protein sequences from cv. B73, because cv. Sugar Bun has not yet been sequenced), and the cRAP database containing common laboratory contaminants (https://www.thegpm.org/crap/index.html). Bacterial species protein sequences were downloaded from NCBI (https://www.ncbi.nlm.nih.gov/bioproject?term=PRJNA357031) and *Z. mays* protein sequences were downloaded from UniProt (https://www.uniprot.org/proteomes/UP000007305).

Given the redundancy in the *Z. mays* genome, we clustered its protein sequences using CD-HIT with a 99% similarity threshold (99). The databases are available in PRIDE (PXD064252).

### Protein identification and data processing

We searched raw MS spectra against the databases described above using the method described in Blakeley-Ruiz *et al.* (100). Briefly, we used the run calibration, SEQUEST HT and percolator nodes in Proteome Discoverer 2.3 (Thermo Fisher Scientific). Search settings can be found in SI Materials and Methods.

We filtered for proteins that were identified with a false discovery rate (FDR) less than 5% and were classified as a Master Protein by Proteome Discoverer. We additionally filtered for proteins that had at least one peptide-spectrum match (PSM) in at least 75% of replicates in at least one condition. We additionally filtered out all *Z. mays* and cRAP proteins, as we were only interested in bacterial proteins for this study. For imputation of missing values in the dataset, we added 1 to all values. We used centered log-ratio transformation prior to statistical analysis. We used the Student’s T-test corrected with the Benjamini-Hochberg false discovery rate (q < 0.05) for multiple hypothesis testing to determine whether protein abundances were significantly different between the *in planta* and *in vitro* conditions.

### Protein annotation

Given the general lack of reliability of functional protein annotations (101), we used a multipronged approach to assign functions to the proteins that were differentially abundant between the two conditions. We first used Mantis to automatically assign functions to all differentially abundant proteins (102). We then manually compared annotations to those generated by NCBI PGAP that were associated with the protein sequences downloaded from NCBI. We additionally used SwissProt, DBCan3, SecRet6, T9GPred, and TCDB to fill in gaps and/or to confirm ambiguous annotations (103–107). The dataset was too large to manually curate all annotations, but we were able to use this method to assign functions to ∼72% (3,997 proteins) of the differentially abundant proteins (Supplementary Data 2). It is important to note that included in the broad categories is a category called “Unknown Function”. This category contained proteins for which we attempted to assign a function but were unable to find a confident match. We then assigned broad and detailed functional categories to the manually annotated, differentially abundant proteins based on the KEGG and MetaCyc pathway databases (108, 109). In total, we assigned 28 broad and 134 detailed functional categories (Supplementary Data 1).

### Hemicellulose degradation assay

To determine whether HRO and CPU are capable of degrading hemicellulose *in vitro*, we conducted a hemicellulose degradation assay. Recipes for the growth media used in this assay can be found in SI Materials and Methods. CPU and HRO were cultured and washed as described above. 100 µL of washed cells were inoculated in 5 mL of each liquid media and placed in a shaker at 30°C at 180 rpm. OD600 was measured at 8 hours and 24 hours. Cultures were pelleted after 24 hours and pellets were frozen at –80°C before proteins were extracted and peptides prepared using S-Trap. Peptides were analyzed by 1D-LC-MS/MS using the protocol described for ELU and BPI in SI Materials and Methods.

### *tssF* knockout mutant generation and growth curves

The methods used to generate the *tssF* mutants can be found in SI Materials and Methods. To determine whether the mutants showed general fitness loss compared to the wildtype, we conducted growth curves of each mutant and wildtype HRO (SI Materials and Methods).

### Plant colonization experiment with *tssF* knockout mutants

The method used for determining whether knocking out *tssF* from HRO would result in a change in plant colonization can be found in SI Materials and Methods.

### Phosphate solubilization assay

We streaked bacterial species from glycerol stocks on their respective selective agar plates and incubated the plates at 30°C for three days. A loopful of bacterial growth was streaked in a single line on Pikovskaya agar plates (HiMedia) and the plates were incubated at 27°C for five days. We then measured the zone of clearance from the edge of the streak to the edge of the cleared zone.

## Declarations

### Ethics approval and consent to participate

Not applicable.

### Consent for publication

Not applicable.

### Availability of data and materials

The mass spectrometry proteomics data and the databases used for protein identification have been deposited to the ProteomeXchange Consortium via the PRIDE (98) partner repository with the dataset identifier PXD064252. The seven bacterial species used in this study are available from DSMZ with the accession numbers 114483, 114565, 114566, 114484, 114485, 114508, 114486.

### Authors’ contributions

A.G.: Experimental design, data collection, data processing, data analysis and interpretation, writing the manuscript

J.C.: T6SS mutant design and construction, T6SS mutant growth curve

N.C.: T6SS mutant design

G.P.: Phosphate solubilization assay

A.N.S.: Conceptual and experimental design of T6SS follow-up experiment, editing

M.R.W.: Conceptualization of the study, editing

A.E.B.: Experimental design, editing

M.K.: Conceptualization of the study, experimental design, mentoring in data analysis and interpretation, writing, editing

*All authors read and approved the final manuscript*.

## Acknowledgements

We made all LC-MS/MS measurements in the Molecular Education, Technology, and Research Innovation Centre (METRIC) at NC State University. We thank the staff of the NC State University Phytotron for the use of their facilities. We thank Dr. Alfredo Blakeley-Ruiz for his guidance on protein annotation and all Kleiner lab members for discussions on methods, results, and data analysis.

