## Supplementary Information for "Differential proteomics of bacteria grown *in vitro* and *in planta* reveals functions used during growth on maize roots"

#### SI Figures and Tables

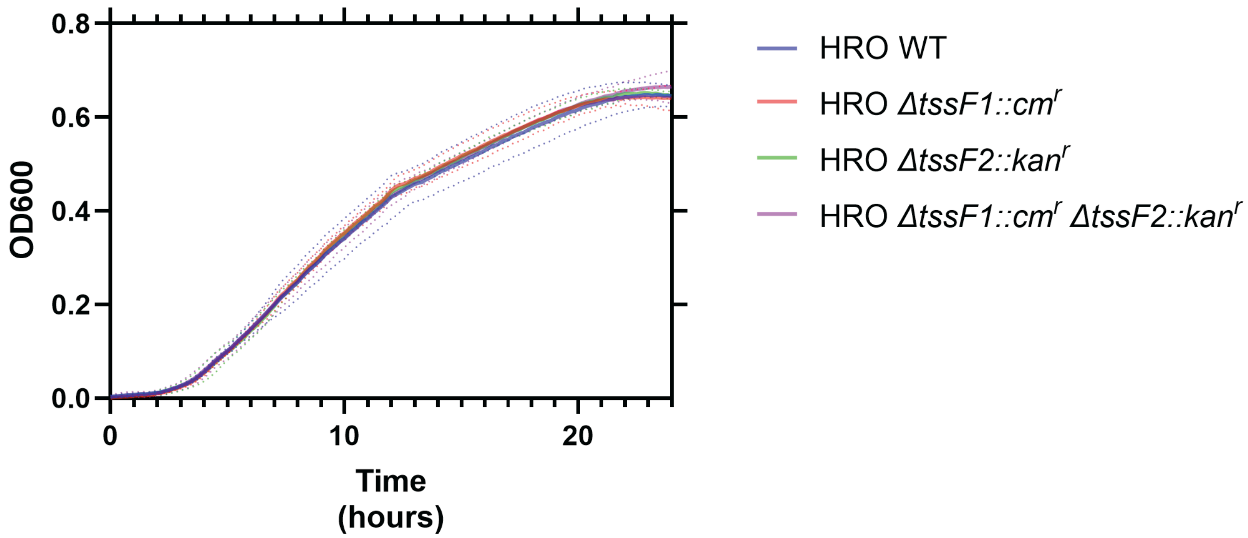

**Figure S1.** Growth curves of HRO wildtype and *tssF* mutants in tryptic soy broth (TSB) over 24 hours, as measured by absorbance at 600 nm. Solid lines indicate mean absorbance for each sample (n = 3), while the dotted lines indicate standard deviation from the mean.

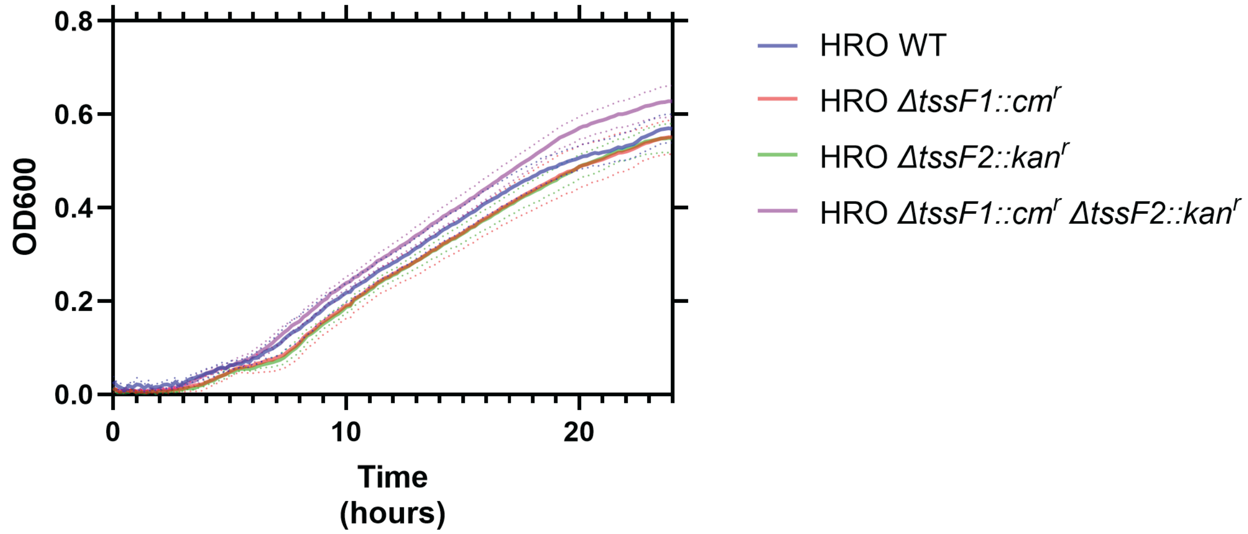

**Figure S2.** Growth curves of HRO wildtype and *tssF* mutants in the defined minimal medium over 24 hours, as measured by absorbance at 600 nm. Solid lines indicate mean absorbance for each sample (n = 8), while the dotted lines indicate standard deviation from the mean.

**Table S1.** Strains, plasmids, and oligonucleotides used to generate *tssF* knockout mutants.

| | **Relevant Characteristics** | **Source or Reference** |
|---|---|---|
| **Strains** | | |
| ***E. coli*** | | |
| DH5α | | NEB Cat# C2987H |
| ***H. robiniae*** | | |
| AA6 | Wild type maize root isolate | (Niu *et al.*, 2017) (1) |
| Δ*tssF1::cm*$^r$ | *TssF1* knockout with *Cm*$^r$ cassette replacement | This study |
| Δ*tssF2*::*kan*$^r$ | TssF2 knockout with *Kan*$^r$ cassette replacement | This study |
| Δ*tssF1::cm*$^r$, Δ*tssF2*::*kan*$^r$ | Double knockout containing both mutations | This study |
| **Plasmids** | | |
| MRTK-clo3' | Knockout vector backbone; ColE1, Amp$^r$ | (van Schaik *et al.*, 2023) (2) |
| MRTK-clo5 | Chloramphenicol resistance cassette; R6kγ, Cm$^r$ | (van Schaik *et al.*, 2023) (2) |
| MRTK-clo6 | Kanamycin resistance cassette; R6kγ, Kan$^r$ | (van Schaik *et al.*, 2023) (2) |
| pJC110 | *TssF1* knockout vector; ColE1, Amp$^r$, Cm$^r$ | This study |
| pJC134 | *TssF2* knockout vector; ColE1, Kan$^r$ | This study |
| **Oligonucleotides** | | |
| JBC_PR_020 | CGAGCCTTCGATCTTGCCTTACGCCGCAG CCGAACGACCGAGC | This study |
| JBC_PR_021 | AACGACGCAAACCATCAATGAGCGGTATCA GCTCACTCAAAGGCGGTAATACGG | This study |
| JBC_PR_022 | TTGAGTGAGCTGATACCGCTCATTGATGGT TTGCGTCGTTCCGGC | This study |

| | | |
|---|---|---|
| JBC_PR_023 | AACCTCTTACGTGCCGATCAAGGAAGACCTGCTTGCCCGTGGC | This study |
| JBC_PR_024 | ACGGGCAAGCAGGTCTTCCTTGATCGGCACGTAAGAGGTTCCAACTTT | This study |
| JBC_PR_025 | ATCACAACAAGCCACCCGACGGCTCACCTTCGGGTGGGCCT | This study |
| JBC_PR_026 | GGCCCACCCGAAGGTGAGCCGTCGGGTGGCTTGTTGTGATGC | This study |
| JBC_PR_027 | CGGTCGTTCGGCTGCGGCGTAAGGCAAGATCGAAGGCTCGACCA | This study |
| JBC_PR_028 | GCAGAAAAAAAGGATCTCAAAAGGCAAGATCGAAGGCTCGAC | This study |
| JBC_PR_029 | GGGGGCGGAGCCTATGGAAACATTGATGGTTTGCGTCGTTCC | This study |
| JBC_PR_030 | AACGACGCAAACCATCAATGTTTCCATAGGCTCCGCCCCCCT | This study |
| JBC_PR_031 | CGAGCCTTCGATCTTGCCTTTTGAGATCCTTTTTTTCTGCGCGTAATC | This study |
| JBC_PR_032 | TGGGCTGGGACAGCTTCATTGCTCGCCAGTCGATTGGCTGAGC | This study |
| JBC_PR_033 | TAGCCGACTTGGGCTGCAGGGATCCAGCATATGCGGTGTGAAA | This study |
| JBC_PR_034 | CACACCGCATATGCTGGATCCCTGCAGCCCAAGTCGGCTACCGCC | This study |
| JBC_PR_035 | GGCCCACCCGAAGGTGAGCCTTAGACTCCTATCCGGGTCGCGCG | This study |
| JBC_PR_036 | CGACCCGGATAGGAGTCTAAGGCTCACCTTCGGGTGGGCCTTTC | This study |
| JBC_PR_037 | AGCGGCGAACAGCCGCTCGAAAAGCCACGTTGTGTCTCAAAAT | This study |
| JBC_PR_038 | TTGAGACACAACGTGGCTTTTCGAGCGGCTGTTCGCCGCTCCCCA | This study |
| JBC_PR_039 | CAGCCAATCGACTGGCGAGCAATGAAGCTGTCCCAGCCCAGCCG | This study |
| JBC_PR_009A | CAGCTTGGGCCGGATGCGCAGGTTCT | This study |
| JBC_PR_010A | CAAGGTCAGGGTCGGCTGACGCCGCC | This study |
| JBC_PR_011A | CTCTCCCTTGAATGCGACTTTCA | This study |
| JBC_PR_012A | TGATCCTCGAAGGCCAAAACCA | This study |
| JBC_PR_013A | CTGGGTGAGTTTCACCAGTTTTG | This study |
| JBC_PR_014A | CAATCGACGTCTATCTTCAAAT | This study |
| JBC_PR_015A | CCATCTTCTTGTCTTCTCTTGT | This study |
| JBC_PR_016A | GGTGGACATCCTCGTGCGCT | This study |
| JBC_PR_017A | GGCTACCGCAACATCCAGGAGTT | This study |

| | | |
|---|---|---|
| JBC_PR_018A | AGCATGAAGCTGTCGGGGCGGATC | This study |
| JBC_PR_019A | AGGTGGACATCCTCGTGCGCTCG | This study |
| JBC_PR_020A | GTTTGCTGTAGCCGTTCAGGCGG | This study |
| JBC_PR_021A | GCTGTCCCGGCCGACGCTAAACA | This study |
| JBC_PR_022A | GATCAACCTGTTTCGCCAGC | This study |
| JBC_PR_023A | CAGCAGCAGCGGCAGGAATTTGT | This study |
| JBC_PR_024A | ATTTCTTTCCAGACTTGTTCAAC | This study |
| JBC_PR_025A | CGGTAACTATCGTCTTGAGTCCA | This study |

### SI Materials and Methods

#### Culturing and inoculation of seven bacterial species for *in vitro* and *in planta* characterization

We cultured all isolates as described by Salvato *et al.* (3). Briefly, we streaked bacterial species from glycerol stocks on selective 0.1x tryptic soy agar (TSA) plates and incubated at 30°C for 48 hours as described by Niu *et al*. (4). An individual colony of each species was inoculated into 5 mL of tryptic soy broth (TSB) and shaken at 30°C for 8 hours. 1 mL of the culture was inoculated into 100 mL of TSB and grown in an incubated shaker at 180 rpm at 30°C overnight. 40 mL of each overnight culture was pelleted by centrifugation at 8000 xg for 8 minutes, re-suspended in 30 mL of phosphate-buffered saline (PBS, VWR International), and pelleted again. The final pellets were re-suspended in 30 mL of PBS buffer.

#### Medium 1: Broth used for the *in vitro* cultivation of BPI, CIN, ELU, HRO, SMA, and PPU in the differential proteomics experiment

Medium components and concentrations are listed in Table S2. Briefly, 0.5x Murashige-Skoog was prepared and autoclaved at 121°C for 30 minutes. Stock solutions of amino acids and vitamins were prepared ahead of time; vitamins were stored at 4°C in the dark and amino acids were stored at -20°C. Stock solutions of malate and glucose were prepared fresh. Malate was pH-adjusted to 7.0 with sodium hydroxide. Glucose, malate, amino acids, and vitamins were filter-sterilized through 0.22 μm filters before adding to sterile, room temperature 0.5x Murashige-Skoog.

**Table S2.** Components of defined, minimal medium used for BPI, CIN, ELU, HRO, PPU, and SMA

| Component | Name | Final Concentration | Brand & Cat. No. |
|---|---|---|---|
| Base Medium | 0.5x Murashige & Skoog | 0.5x | RPI; M10200 |
| Carbon Source | Malate | 1% w/v | |
| Carbon Source | Glucose | 1% w/v | |
| Amino Acid | Arginine | 50 μg/mL | |
| Amino Acid | Proline | 50 μg/mL | |
| Amino Acid | Serine | 50 μg/mL | |
| Amino Acid | Asparagine | 50 μg/mL | |
| Amino Acid | Leucine | 50 μg/mL | |
| Amino Acid | Isoleucine | 50 μg/mL | |
| Amino Acid | Lysine | 50 μg/mL | |
| Amino Acid | Cysteine | 50 μg/mL | |
| Amino Acid | Glutamine | 50 μg/mL | |
| Vitamin | Riboflavin (B2) | 0.2 μg/mL | |
| Vitamin | Nicotinic Acid (B3) | 1 μg/mL | |
| Vitamin | Cobalamin (B12) | 0.02 μg/mL | |

**Medium 2: Broth used for the *in vitro* cultivation of CPU in the differential proteomics experiment**

Defined, minimal medium used for the *in vitro* growth of the CPU in the differential proteomics experiment. This medium was prepared using the same method as Medium 1, but with the addition of pyridoxine (vitamin B4), pantothenate (vitamin B5), and biotin (vitamin B7) (Table S3), as we previously found in Garrell *et al.*(5) that CPU required these three additional vitamins for growth, while the other six SynCom species did not.

**Table S3.** Components of defined, minimal medium used for CPU.

| Component | Name | Final Concentration | Brand & Cat. No. |
|---|---|---|---|
| | Medium 1 | | |
| Vitamin | Pyridoxine (B4) | 1 µg/mL | |
| Vitamin | Pantothenate (B5) | 1 µg/mL | |
| Vitamin | Biotin (B7) | 0.01 µg/mL | |

**Medium 3: Broth used as the "No Carbon" medium in the hemicellulose degradation assay**

While this medium does contain carbon in the form of amino acids, there is no additional, primary carbon source. This medium was prepared using the same method and components as Medium 2, but glucose and malate were omitted from this medium.

**Medium 4: Broth used as the "Xylan" medium in the hemicellulose degradation assay**

This medium was prepared using the same method and components as Medium 2, but glucose and malate were omitted from this medium and replaced with filter-sterilized 1% w/v xylan from corn core (TCI America).

**Protein extraction and peptide preparation**

HRO, SMA, PPU, CIN, and CPU peptide samples were analyzed by one-dimension LC-MS/MS as described by Mordant and Kleiner (6): "For each sample, [1200] ng of tryptic peptides were loaded with an UltiMate 3000 RSLCnano liquid chromatograph (Thermo Fisher Scientific) in loading solvent A (2% acetonitrile, 0.05%

trifluoroacetic acid) onto a 5-mm, 30-µm-inner diameter C18 Acclaim PepMap100 precolumn and desalted (Thermo Fisher Scientific). Peptides were then separated on a 75-cm × 75-µm analytical EASY-Spray column packed with PepMap RSLC C18, 2-µm material (Thermo Fisher Scientific) heated to 60°C via the integrated column heater at a flow rate of 300 nL $min^{-1}$ using a 140 min gradient going from 95% buffer A (0.1% formic acid) to 31% buffer B (0.1% formic acid, 80% acetonitrile) in 102 min, then to 50% B in 18 min, to 99% B in 1 min, and ending with 99% B. Carryover was reduced by wash runs (injection of 20 µL acetonitrile with 99% eluent buffer B) between samples.

"The analytical column was connected to a Q Exactive HF hybrid quadrupole-Orbitrap mass spectrometer (Thermo Fisher Scientific) via an EASY-Spray source. Eluting peptides were ionized via electrospray ionization (ESI). MS1 spectra were acquired by performing a full MS scan at a resolution of 60,000 on a 380 to 1,600 m/z window. MS2 spectra were acquired using a data-dependent approach by selecting for fragmentation the 15 most abundant ions from the precursor MS1 spectra. A normalized collision energy of 25 was applied in the high cell density (HCD) cell to generate the peptide fragments for MS2 spectra. Other settings of the data-dependent acquisition included a maximum injection time of 100 ms, a dynamic exclusion of 25 s, and exclusion of ions of +1 charge state from fragmentation. About 60,000 MS/MS spectra were acquired per sample."

ELU and BPI samples were analyzed using a similar method to that above, but with a few modifications. For each sample, 1000 ng of tryptic peptides were loaded using the method and liquid chromatograph described above. The analytical column was connected via an EASY-Spray source to an Exploris 480 hybrid quadrupole-Orbitrap mass spectrometer (Thermo Fisher Scientific). Peptides were separated on the analytical column using the same 140 min gradient as above. MS1 spectra were obtained with a full MS scan at a resolution of 60,000 on a 380 to 1,600 m/z window. MS2 spectra were generated by choosing the 15 most abundant peptides from the precursor MS1 spectra for fragmentation. Fragmentation was performed in the ion routing multipole with a normalized collision energy of 27%. For MS2 we used a maximum injection time of 50 ms, a dynamic exclusion of 25 s, and exclusion of ions of +1 charge state from fragmentation. Roughly 100,000 MS/MS spectra were acquired per sample.

#### **Protein identification and data processing**

We searched raw MS spectra against the databases described above using the method described in Blakeley-Ruiz *et al.* (7). Briefly, we used the run calibration, SEQUEST HT and percolator nodes in Proteome Discoverer 2.3 (Thermo Fisher Scientific). We used the following search settings: trypsin (full), 2 missed cleavages, 10 ppm precursor mass tolerance, 0.1 Da fragment mass tolerance. We included the following dynamic modifications: oxidation on M (+15.995 Da), deamidation on N, Q, R (0.984 Da), and acetyl on the protein N terminus (+42.011 Da). We also included the static modification carbamidomethyl on C (+57.021 Da).

#### ***tssF* knockout mutant generation and growth curves**

To generate the *tssF* mutants, we constructed two integrative vectors using the pMRMTK-clo3' plasmid backbone (Addgene plasmid # 203924;

http://n2t.net/addgene:203924; RRID:Addgene_203924) generated in a previous study (2). The *tssF1* integrative vector (pJC110) was assembled from the backbone, a chloramphenicol resistance (*cm*$^r$) cassette, and 1 kb genomic regions flanking the *tssF1* site in *Herbaspirillum robiniae* AA6 (HRO). The *tssF2* integrative vector (pJC134) was similarly constructed using a kanamycin resistance (*kan*$^r$) cassette and the 1 kb *tssF2*-flanking regions. DNA fragments were amplified by PCR using the appropriate primers (Table S1), followed by DpnI digestion, gel extraction and purification. Gibson assembly for each integrative vector was performed using the NEBuilder® HiFi DNA Assembly Master Mix (Cat #: E2621L) following the protocol for a 4-part assembly. Initial transformations were performed in NEB® 5-alpha Competent *Escherichia coli* (High Efficiency) (Catalog #: C2987H), and constructs were verified by Plasmidsaurus using Oxford Nanopore Technology with custom analysis and annotation. To generate the single knockout mutants, electrocompetent HRO cells were prepared (as previously described in Van Schaik *et al.* (2)) and 50 µL of the cell suspension was transformed with 1192 ng of pJC110 (resulting in a Δ*tssF1::cm*$^r$ knockout mutant) or 724 ng of pJC134 (resulting in a Δ*tssF2*::*kan*$^r$ knockout mutant). To generate the double knockout mutant, HRO Δ*tssF1::cm*$^r$ was transformed with 484 ng of pJC134, resulting in a Δ*tssF1::cm*$^r$*,* Δ*tssF2*::*kan*$^r$ mutant (2). Transformants were plated on the appropriate selective media (LB + Cm or LB + Kan). As pJC110 contained an AmpR cassette, the Δ*tssF1::cm*$^r$ transformants were plated on LB + Amp to counterselect against ampicillin resistance. Integration of antibiotic resistance cassettes to replace *tssF1* and *tssF2* was confirmed by PCR and amplicon sequencing.

To determine whether the mutants showed any general loss in fitness compared to the wildtype, we conducted growth curves of each mutant and wildtype HRO over 24 hours. Cells were plated from frozen stock onto full strength TSA containing the appropriate antibiotics for each mutant. Three colonies were chosen as biological replicates and grown in TSB containing the appropriate antibiotics for each mutant overnight in a 30°C shaking incubator at 250 rpm. Cell cultures were diluted to an OD600 of 0.01, and the 3 biological replicates for each condition were aliquoted into a 96-well plate with TSB along with an uninoculated control. Absorbance at 600 nm was measured over a 24-hour period using the Tecan Sunrise microplate reader with continuous shaking (250 rpm) at 30°C. Growth curve graphs were generated in GraphPad Prism 10.

**Plant colonization experiment with *tssF* knockout mutants**

Seeds for the plant colonization experiment were pre-germinated 2 days before experiment set up. We sterilized and rinsed *Zea mays* cv. Sugar Bun seeds (untreated; Johnny's Selected Seeds) using the method described above for *in planta* characterization. We placed strips of autoclaved germination paper (Fisher Scientific Cat. No. NC1466201) into the bottoms of sterile 24-well plate wells, then placed a sterilized seed into each well. We then added 200 µL sterile $dH_2O$ to each seed and covered the plates. Plates were incubated at 30°C in the dark for 48 hours.

We inoculated the wildtype HRO strain and its mutants on their respective selective plates (see Medium 5-8 described below) and incubated them at 30°C for 48 hours. An individual colony of each species was inoculated into 5 mL of TSB with 50 µg/mL kanamycin (Fisher Scientific CAS No. 25389-94-0) and/or chloramphenicol (Millipore Sigma CAS No. 56-76-7) for the knockout mutants and shaken at 180 rpm at 30°C for 8 hours. 500 µL of the culture was inoculated into 50 mL of tryptic soy broth (with 50 µg/mL kanamycin and/or chloramphenicol for the knockout mutants) and grown in an incubated shaker at 180 rpm at 30°C overnight. Cells were collected, washed, and diluted in the same manner as described in the culturing and inoculation section above.

The seedlings that were germinated in 24-well plates were planted and inoculated in sterile WhirlPak bags following the protocol described above. Plants were placed in a growth chamber on 12 hour cycles of fluorescent light and dark at 27°C and 23°C, respectively, for 14 days. 10 bags, each containing one seed, were inoculated for each of the conditions.

Roots were collected, rinsed, and cut following the method described above. We note that despite pre-germination, not all plants grew after transplantation to the WhirlPak bags. We therefore proceeded with 8-9 replicates. 0.3 g of root fragments were placed into 1 mL sterile PBS with six sterile 3 mm glass beads, then vortexed 3 times for 1 minute, with 10 seconds dwell time between each vortex. Following the protocol outlined by Niu and Kolter, 20 µL of the slurry was then serially diluted in a 96-well plate for $10^{-1}$ to $10^{-8}$ dilutions (4). 10 µL of each dilution was then spotted on a selective plate using a multichannel pipette, and plates were tilted to spread the drops. Plates were incubated at 30°C for 36 hours, before colony forming units were counted.

**Medium 5: Agar used for cultivation of wildtype HRO for plant colonization experiment with *tssF* knockout mutants**

This medium was prepared following the protocol outlined in Niu *et al.* (4). Briefly, 1.38 g of tryptic soy broth without dextrose (TSB), 7.5 g agar, and 500 mL distilled water were mixed and autoclaved at 121°C for 30 min. Once cooled to approx. 60°C, added 1,000 µL nalidixic acid (stock solution at 5 mg/mL), 200 µL colistin (stock solution at 10 mg/mL), and 1,000 µL lincomycin (stock solution at 50 mg/mL).

**Medium 6: Agar used for cultivation of HRO Δ*tssF1::cm*$^r$ knockout mutant for plant colonization experiment with *tssF* knockout mutants**

Agar was prepared following the method used for Medium 5, with the addition of 500 µL chloramphenicol (stock solution at 34 mg/mL).

**Medium 7: Agar used for cultivation of HRO Δ*tssF2*::*kan*$^r$ knockout mutant for plant colonization experiment with *tssF* knockout mutants**

Agar was prepared following the method used for Medium 5, with the addition of 500 µL kanamycin (stock solution at 50 mg/mL).

**Medium 8: Agar used for cultivation of HRO Δ*tssF1::cm*$^r$, Δ*tssF2*::*kan*$^r$ mutant for plant colonization experiment with *tssF* knockout mutants**

Agar was prepared following the method used for Medium 5, with the addition of 500 µL chloramphenicol (stock solution at 34 mg/mL) and 500 µL kanamycin (stock solution at 50 mg/mL).

alters gut microbiota composition and function. ISME J wraf048.
